# Tumor Cytokine-Induced Hepatic Gluconeogenesis Contributes to Cancer Cachexia: Insights from Full Body Single Nuclei Sequencing

**DOI:** 10.1101/2023.05.15.540823

**Authors:** Ying Liu, Ezequiel Dantas, Miriam Ferrer, Yifang Liu, Aram Comjean, Emma E. Davidson, Yanhui Hu, Marcus D. Goncalves, Tobias Janowitz, Norbert Perrimon

## Abstract

**Summary:** A primary cause of death in cancer patients is cachexia, a wasting syndrome attributed to tumor-induced metabolic dysregulation. Despite the major impact of cachexia on the treatment, quality of life, and survival of cancer patients, relatively little is known about the underlying pathogenic mechanisms. Hyperglycemia detected in glucose tolerance test is one of the earliest metabolic abnormalities observed in cancer patients; however, the pathogenesis by which tumors influence blood sugar levels remains poorly understood. Here, utilizing a *Drosophila* model, we demonstrate that the tumor secreted interleukin-like cytokine Upd3 induces fat body expression of *Pepck1* and *Pdk*, two key regulatory enzymes of gluconeogenesis, contributing to hyperglycemia. Our data further indicate a conserved regulation of these genes by IL-6/JAK STAT signaling in mouse models. Importantly, in both fly and mouse cancer cachexia models, elevated gluconeogenesis gene levels are associated with poor prognosis. Altogether, our study uncovers a conserved role of Upd3/IL-6/JAK-STAT signaling in inducing tumor-associated hyperglycemia, which provides insights into the pathogenesis of IL-6 signaling in cancer cachexia.

**Graphical Abstract:** 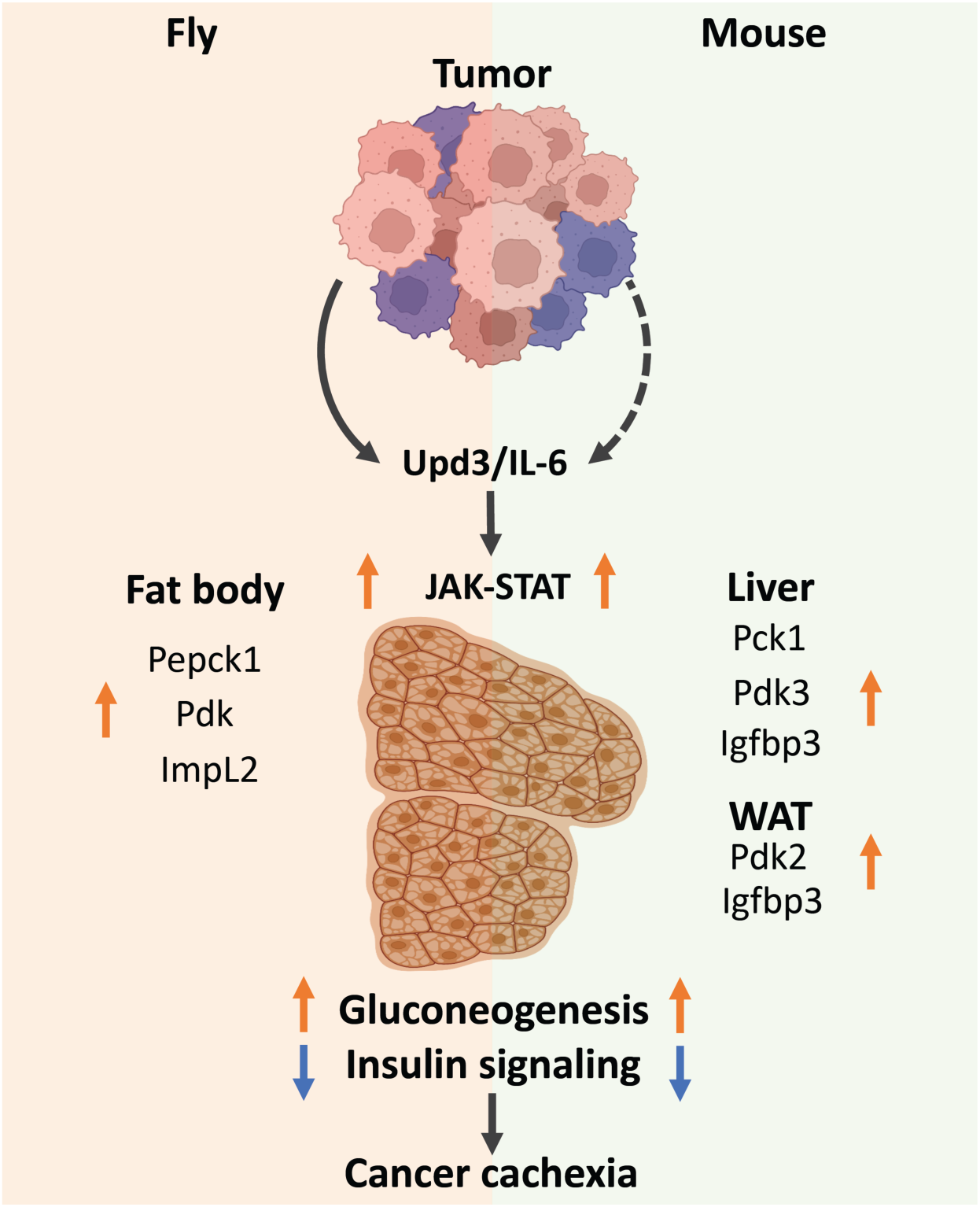

## Introduction

More than 40% of cancer patients suffer from cachexia, a tumor-driven life-threatening condition with symptoms of massive weight loss, general inflammation, weakness, and fatigue (Dewys et al., 1980; Teunissen et al., 2007; Argilés et al., 2010). A prominent driving force of these cachectic symptoms is the metabolic dysregulation stimulated by tumors, such as the systemic reprogramming of glucose metabolism (Warburg, 1956; Holroyde et al., 1975; Hanahan and Weinberg, 2011; Petruzzelli and Wagner, 2016). In fact, higher than normal blood glucose levels (hyperglycemia) as revealed by glucose tolerance testing is the earliest metabolic abnormality observed in cancer patients, and has been previously associated with insulin resistance (Rohdenburg et al., 1919; Jasani et al., 1978; Lundholm et al., 1978; Tayek, 1992; Fearon et al., 2012). As insulin signaling is required for retaining glucose intake and glycolysis in peripheral tissues, cancer patients with reduced insulin sensitivity may have a declined glucose degradation rate, leading to hyperglycemia (Wu et al., 2005; Guo et al., 2012). In support of this model, CD2F1 mice with colon-26 adenocarcinoma tumors show blunted blood glucose response to insulin and reduced phosphorylation of Akt in muscle and adipose tissues (Asp et al., 2010). Despite these observations, our understanding of how tumors induce insulin resistance in host organs remains incomplete. Possible candidates are the tumor-induced expression of IGF-binding proteins (IGFBP1-6), which can antagonize insulin/IGF signaling (Baxter, 2014; Remsing Rix et al., 2022). Studies in cancer patients have proposed a role for IGFBP2 and IGFBP3 in cachexia (Huang et al., 2016; Dong et al., 2021), although whether IGFBPs induce hyperglycemia in cancer patients has not been reported. Interestingly, in two *Drosophila* models of organ wasting/cachexia, ecdysone-inducible gene L2 (ImpL2), an insulin binding protein that is functionally equivalent to mammalian IGFBPs (Honegger et al., 2008), is secreted from either gut tumors (Kwon et al., 2015) or tumorous imaginal discs (Figueroa Clarevega and Bilder, 2015), and causes hyperglycemia by repressing systemic insulin signaling. Collectively, these studies suggest a conserved mechanism of IGFBPs/ImpL2 in tumor-induced systemic insulin resistance.

Elevated hepatic glucose production (gluconeogenesis) can also result in hyperglycemia (Bock et al., 2007; Meshkani and Adeli, 2009; Petersen and Shulman, 2018). Gluconeogenesis requires a number of enzymes such as phosphoenolpyruvate carboxykinase (PEPCK), pyruvate carboxylase (PC), fructose 1,6-bisphosphatase (FBP), and glucose 6-phosphatase (G6P) to synthesize glucose from oxaloacetate through multiple reactions (Melkonian et al., 2022). Among them, PEPCK is a rate-limiting enzyme that catalyzes the first reaction of gluconeogenesis, converting oxaloacetate into phosphoenolpyruvate toward glucose production (Rognstad, 1979; Yu et al., 2021). In addition to PEPCK, Pyruvate dehydrogenase kinase (PDK) is an important regulator of gluconeogenesis (Tao et al., 2013; Herbst et al., 2014; Zhang et al., 2014). In the liver, PDK inhibits the conversion of pyruvate to acetyl-CoA through inactivation of the pyruvate dehydrogenase complex (PDC), which redirects pyruvate toward gluconeogenesis (Zhang et al., 2014; Melkonian et al., 2022). In other organs that do not undergo gluconeogenesis, PDK reduces the levels of acetyl-CoA, leading to the repression of fatty acid synthesis from glucose (Huang et al., 2002; Zhang et al., 2014). Notably, tumor-bearing mice and rats with cachexia display increased hepatic expression of PEPCK (Tisdale, 2009; Narsale et al., 2015; Viana et al., 2018), suggesting increased hepatic gluconeogenesis in these animals. Indeed, the association between increased hepatic gluconeogenesis activity and cancer cachexia has been observed for decades (Holroyde et al., 1975, 1984). Importantly, a physiological characteristic of gluconeogenesis is that it does not yield but consume energy (Blackman, 1982). Thus, elevated levels of hepatic gluconeogenesis in cancer patients may contribute to their energy and weight loss (Stein, 1978; Bongaerts et al., 2006). Despite these observations, how hepatic gluconeogenesis activation in cancer patients is stimulated remains poorly understood. One model is that gluconeogenesis is activated by metabolic substrates derived from tumors, such as lactate, alanine, or glycerol (Waterhouse, 1974; Stein, 1978). It has also been proposed that reduced insulin signaling activity may promote gluconeogenesis (Yoshikawa et al., 2001; Agustsson et al., 2011; Winter et al., 2012). Further, inflammatory factors have also been proposed to play a role in the progression of insulin resistance (Yoshikawa et al., 2001). For example, administration of the anti-inflammatory thiol compound Pyrrolidine dithiocarbamate (PDTC) can inhibit aberrant hepatic PEPCK induction in a mouse model of intestinal cancer (Apc^Min/+^) (Narsale et al., 2015), indicating a role for inflammation in tumor-induced gluconeogenesis. A particularly relevant inflammatory cytokine is interleukin-6 (IL-6), which activates the Janus kinase/Signal transducer and activator of transcription (JAK STAT) signaling pathway. IL-6 has been shown to positively correlate with weight loss and mortality in cancer patients (Strassmann et al., 1992; Scott et al., 1996; Barton, 2001; Moses et al., 2009; Argilés et al., 2019). However, whether hepatic gluconeogenesis is directly targeted by inflammatory pathways, or is a secondary effect of systemic inflammation in cancer patients, requires further studies.

In recent years, a number of organ wasting/cachexia models have emerged in *Drosophila* (Bilder et al., 2021; Liu et al., 2022). In particular, expression of an activated form of the Yorkie (Yki)/Yap oncogene in adult intestinal stem cells (ISCs) generates tumors associated with cachectic properties (Kwon et al., 2015). These tumors secrete at least three factors: ImpL2, Pvf1 and Upd3 (Kwon et al., 2015; Song et al., 2019; Ding et al., 2021). By antagonizing insulin signaling, ImpL2 leads to reduced anabolism in peripheral tissues (Kwon et al., 2015; Ding et al., 2021). Pvf1, a cytokine reminiscent of PDGF/VEGF, activates ERK signaling in peripheral tissues, triggering catabolism (Song et al., 2019). Finally, Upd3, the fly ortholog of IL-6, also induces *ImpL2* expression in peripheral tissues, impairing insulin signaling and contributing to body wasting (Ding et al., 2021). Since cancer cachexia is a multi-organ syndrome, understanding it requires knowledge of how various metabolic tissues respond to tumors. Here, we systemically characterize the pathogenesis underlying cancer cachexia in the fly *yki*^act^ gut tumor model using recently established technology of single nuclei RNAseq (snRNAseq) of whole-body flies. By transcriptome profiling of gut tumors and wasting tissues, we uncovered that tumor-induced Upd3/JAK-STAT signaling directly targets and promotes *Pepck1* and *Pdk* expression in the fat body. This is reminiscent of the activation of hepatic gluconeogenesis, as this tissue performs some of the functions of the liver. Importantly, our results suggest that this cachectic role of Upd3/IL-6 is conserved in mouse models of cachexia. Altogether, our findings may pave the way to a new therapeutic strategy of targeting hepatic gluconeogenesis in IL-6 related cancer cachexia.

## Results

### Body-wide gene expression dynamics of Yki flies

Aiming at a comprehensive understanding of tumor induced wasting in host organs, we examined the full body transcriptome of flies with *yki*^act^ gut tumors (*esg>yki*^act^; referred to as Yki flies) at single-nuclei resolution. Yki flies develop tumors at 2 days following *yki*^act^ induction, and tumors encompass most of the gut at Day 4. Wasting of peripheral organs and bloating (accumulation of body fluid) in these animals starts at Day 5 and is severe at Day 8 (Song et al., 2019) (Figure 1AB). To decipher the transcriptional changes occurring in peripheral tissues, we isolated nuclei of Yki flies at Day 5 and Day 8 following tumor induction together with the appropriate controls, then performed single-nuclei RNA-sequencing (snRNA-seq). Note that we did not include heads in these samples as our study was focused on changes occurring in the gut, muscle, fat body/adipose tissues, and oenocytes. In total, 122,898 nuclei were profiled (25,146 from control and 42,375 from Yki flies at Day 5; 19,050 from control and 36,327 from Yki flies at Day 8), and a median of 559 genes and 990 unique molecular identifiers (UMIs) per nuclei were obtained across all conditions. Next, we generated a uniform manifold approximation and projection (UMAP) plot of all cells and identified 31 cell clusters (Figure 1C, Figure S1A). We annotated these clusters based on the marker genes of *Drosophila* cell types reported by the Fly Cell Atlas (Li et al., 2022) (Figure S1AB). These clusters represent cells from all the major organs (Figure 1C) with an average of 12,039 expressed genes per organ (Figure S1C).

**Figure 1.**
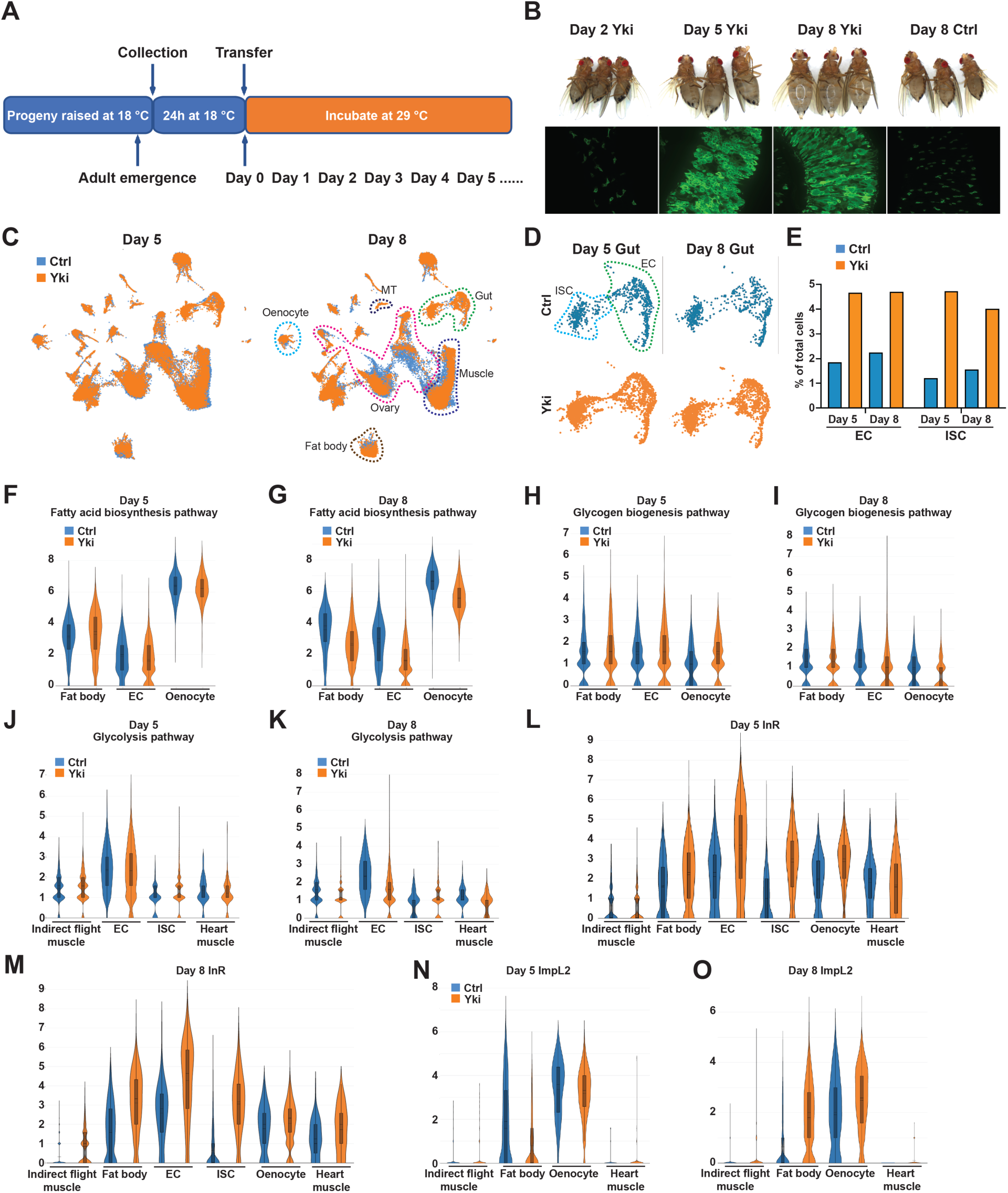
Full-body single-nucleus transcriptome survey of *Drosophila* Yki flies. **(A)** Experimental design of tumor induction in flies. **(B)** Representative gut tumor and phenotypes of Yki flies at Day 2, Day 5, Day 8, and control flies at Day 8. **(C)** UMAP visualization of cell clusters of control (coral) and Yki (indigo) flies at Day 5 and Day 8. **(D)** UMAP visualization of intestinal stem cells (ISC) and enterocyte (EC) clusters of control and Yki flies at Day 5 and 8. **(E)** EC and ISC proportion comparison between control and Yki flies at Day 5 and 8. Expression levels of fatty acid biosynthesis pathway genes at Day 5 **(F)** and Day 8 **(G)** in the fat body, EC, and oenocyte cell clusters represented by violin plots. Expression levels of glycogen biosynthesis pathway genes at Day 5 **(H)** and Day 8 **(I)** in the fat body, EC, and oenocyte cell clusters visualized by violin plots. Expression levels of glycolysis pathway genes at Day 5 **(J)** and Day 8 **(K)** in the indirect flight muscle, EC, ISC, and heart muscle cell clusters visualized by violin plots. Expression levels of *InR* at Day 5 **(L)**, Day 8 **(M),** *ImpL2* at Day 5 **(N)** and Day 8 **(O)** in the indirect flight muscle, fat body, EC, ISC, oenocyte, heart muscle cell clusters visualized by violin plots. See also Figure S1.

Expression of *yki*^act^ in ISCs increases cell proliferation. In line with this, a higher portion of ISC nuclei were recovered in Yki flies compared to control (Yki flies 4.7% vs Control 1.2% at Day 5, 4.0% vs 1.6% at Day 8) (Figure 1DE, Figure S1D). In addition, the portion of EC nuclei was also increased, indicating that *yki*^act^ progenitors are still capable of differentiation (Yki flies 4.7% vs Control 1.8% at Day 5, 4.7% vs 2.3% at Day 8) (Figure 1DE, Figure S1D). Consistent with the previous observation that *yki*^act^ gut tumors lead to ovary atrophy (Kwon et al., 2015), a reduced number of ovarian nuclei were recovered from Yki flies including several germline cell and follicle cell clusters (Figure 1C, Figure S1D). Interestingly, although depletion of triglycerides and glycogen were observed in Yki flies (Kwon et al., 2015), no dramatic changes in the fat body cluster were observed (Figure 1C, Figure S1D). This indicates that the reduction of energy storage in Yki flies is not due to loss of fat body cells and suggests it might instead be an outcome of tumor-induced metabolic reprogramming.

### Reprograming of host metabolism by the tumorous gut

*yki*^act^ gut tumors disturb metabolic homeostasis of host organs, which contributes to host body wasting (Kwon et al., 2015; Song et al., 2019; Ding et al., 2021). To characterize how the metabolism of each organ is affected by the tumorous gut, we assessed the gene expression profile of relevant metabolic pathways (Figure S1E). In Yki flies, the progression of host organ wasting is accompanied with reduced levels of energy storage (lipids and glycogen) (Kwon et al., 2015). The reduction of body lipid storage in Yki flies could result from reduced fatty acid biosynthesis, reduced fatty acid elongation, elevated fatty acid degradation, or a combination of these processes. Thus, we examined the expression of relevant enzymes across the clusters defining the various organs (Figure 1FG, Figure S1F-K). Consistent with a previous observation that the wasting phenotype of Yki flies initiates at Day 5 (Song et al., 2019), expression of fatty acid biosynthesis genes showed minor changes at Day 5 but was significant reduced at Day 8 in the fat body, ECs, and oenocytes (Figure 1FG, Figure S1FG). While changes at Day 5 and in other clusters were minor, fatty acid elongation genes were reduced in hindgut cell clusters at Day 8 (Figure S1HI). In contrast, no dramatic increase of fatty acid degradation genes was observed across all clusters and time points (Figure S1JK). Altogether, these data suggest that the reduction in lipid levels observed in Yki flies results primarily from the inhibition of fatty acid biosynthesis. Next, we analyzed the expression of genes involved in glycogenesis and glycogenolysis (Figure 1HI, Figure S1L-O). The expression levels of glycogen degradation genes were decreased in Yki flies in the fat body cluster at both Day 5 and Day 8, with a dramatic reduction in ECs at Day 8 (Figure S1NO), suggesting an attenuated level of glycogen degradation in Yki flies. Thus, the reduction of glycogen in Yki flies is probably due to abnormalities in glycogenesis. In support of this model, glycogenesis genes displayed reduced expression in ECs and oenocytes at Day 8 (Figure 1I). Notably, we did not observe a reduction of glycogenesis genes in these clusters at Day 5 (Figure 1H), suggesting that the decrease in glycogenesis follows tumor formation. Collectively, these data suggest that the depletion of energy storage in Yki flies is caused by a decrease in systemic anabolic metabolism.

In addition to reduced levels of energy storage, Yki flies display increased amounts of glucose (Kwon et al., 2015), leading us to investigate the expression profile of glycolysis-related genes (Figure 1JK, Figure S1PQ). Glycolysis genes were more evenly expressed in all clusters, likely because of their ubiquitous roles in generating energy in different cell types (Figure S1PQ). At Day 5, Yki flies showed minor expression changes of glycolysis genes (Figure 1J, Figure S1P). Interestingly, a wide inhibition of glycolysis gene expression was apparent at Day 8 in various clusters, including the indirect flight muscle, heart muscle, and ECs (Figure 1K, Figure S1Q). Notably, while most cell clusters displayed reduced or unchanged levels of glycolysis gene expression at Day 8, ISCs, where the tumor is induced, showed elevated glycolysis (Figure 1K, Figure S1Q). This observation suggests that tumor cells can overcome the effects of ImpL2 and sustain energy generation from glucose while overall glycolysis levels are repressed in Yki flies, consistent with the model proposed previously based on bulk RNAseq data (Kwon et al., 2015; Lee et al., 2021).Thus, suppressed glycolysis in peripheral organs may be the origin of the elevated glucose levels in Yki flies.

In mammals, reduction of glycolysis, lipogenesis, and glycogenesis are signs of decreased insulin signaling (Wu et al., 2005; Guo et al., 2012). Indeed, we observed a broad upregulation of *InR* expression in cell clusters of muscle, fat body, ECs, ISCs, and oenocytes, with prominent increases in muscle and fat body from Day 5 to Day 8 (Figure 1LM), indicating reduced insulin signaling in these tissues (Puig and Tjian, 2005). These observations are consistent with the observation that secretion of ImpL2 from both Yki tumors and host organs repress insulin signaling, leading to the systemic decline of glycolysis in Yki flies (Kwon et al., 2015; Ding et al., 2021). In support of this model, we observed upregulation of *ImpL2* in muscle clusters at both Day 5 and 8, and in oenocytes at Day 8, and a prominent increase in *ImpL2* levels in the fat body at Day 8 (Figure 1NO). Notably, although expression of *ImpL2* was increased at Day 5 in a few cell clusters, the systemic decline of glycolysis, lipogenesis, and glycogenesis was only detected later at Day 8 when *ImpL2* was significant upregulated in the fat body (Figure 1NO, Figure S1RS), suggesting a significant effect of fat body *ImpL2* expression on whole-body insulin signaling levels. Altogether, our data suggests that tumor-induced expression of *ImpL2* in hepatocytes and adipocytes contributes significantly to cancer cachexia-related insulin resistance.

### Fat body gluconeogenesis contributes to elevated trehalose levels in Yki flies

Another prominent metabolic feature of Yki flies is elevated levels of trehalose (Kwon et al., 2015), the major insect “blood sugar” (Matsuda et al., 2015; Yoshida et al., 2016). In animals, blood sugar homeostasis is maintained through multiple mechanisms, including gluconeogenesis (Hatting et al., 2018). Thus, our observation of increased trehalose levels in Yki flies suggests a potential elevation of gluconeogenesis. In mammals, Glucose-6 Phosphatase (G6p) catalyzes the last step of gluconeogenesis which produces glucose, whereas in flies the glucose 6-phosphate is catalyzed by Trehalose-6-phosphate synthase 1 (Tps1) that generates trehalose (Matsuda et al., 2015; Yoshida et al., 2016) (Figure 2A). Interestingly, we observed abundant expression of *Tps1* in the fat body and malpighian tubules (MT), suggesting that gluconeogenesis occurs mostly in these tissues (Figure S2AB). Next, we analyzed the expression of gluconeogenesis genes (Figure S1E) which, while reduced or not changed in most cell clusters, were significant increased in the fat body at Day 8 (Figure 2BC, Figure S2CD), indicative of elevated trehalose production. Interestingly, *Pepck1* and *Pdk*, two decisive regulators that promote gluconeogenesis (Huang et al., 2002; Zhang et al., 2014; Yu et al., 2021), were significantly increased at Day 8 in the fat body (Figure 2DE, Figure S2EF). Thus, we hypothesized that the increased expression of *Pepck1* and *Pdk* in fat body of Yki flies leads to elevated gluconeogenesis, contributing to the hyperglycemic phenotype. Consistent with this model, a significant increase of whole-body glucose and trehalose levels in Yki flies was detected at Day 8, when the expression levels of *Pepck1* and *Pdk* were elevated prominently, but not at Day 5 (Figure 2FG). To further investigate the role of fat body Pepck1 and Pdk in controlling body carbohydrates levels, we decreased *Pepck1* in the fat body of Yki flies using two dual-binary-systems, GAL4/UAS and LexA/LexAop, to manipulate gene expression in the gut and in the fat body, respectively (Saavedra et al., 2021). *Pepck1* depletion in the fat body of Yki flies showed no obvious change of the GFP-labeled gut tumor cells but suppressed the bloating phenotype (Figure 2H). In addition, whole-body trehalose levels were significantly reduced in these flies (Figure 2J, Figure S2H). Downregulation of *Pdk* had a similar effect as downregulation of *Pepck1*: gut tumors were not obviously changed, bloating was suppressed, and trehalose levels were decreased (Figure 2KM, Figure S2MQ). Interestingly, glucose and other analyzed metabolites were not affected in *Pepck1* or *Pdk* depleted flies (Figure 2I, L, Figure S2G, I-L, N-P). Altogether, these findings suggest that fat body gluconeogenesis induced by *Pepck1* and *Pdk* upregulation leads to elevated trehalose levels in Yki flies.

**Figure 2.**
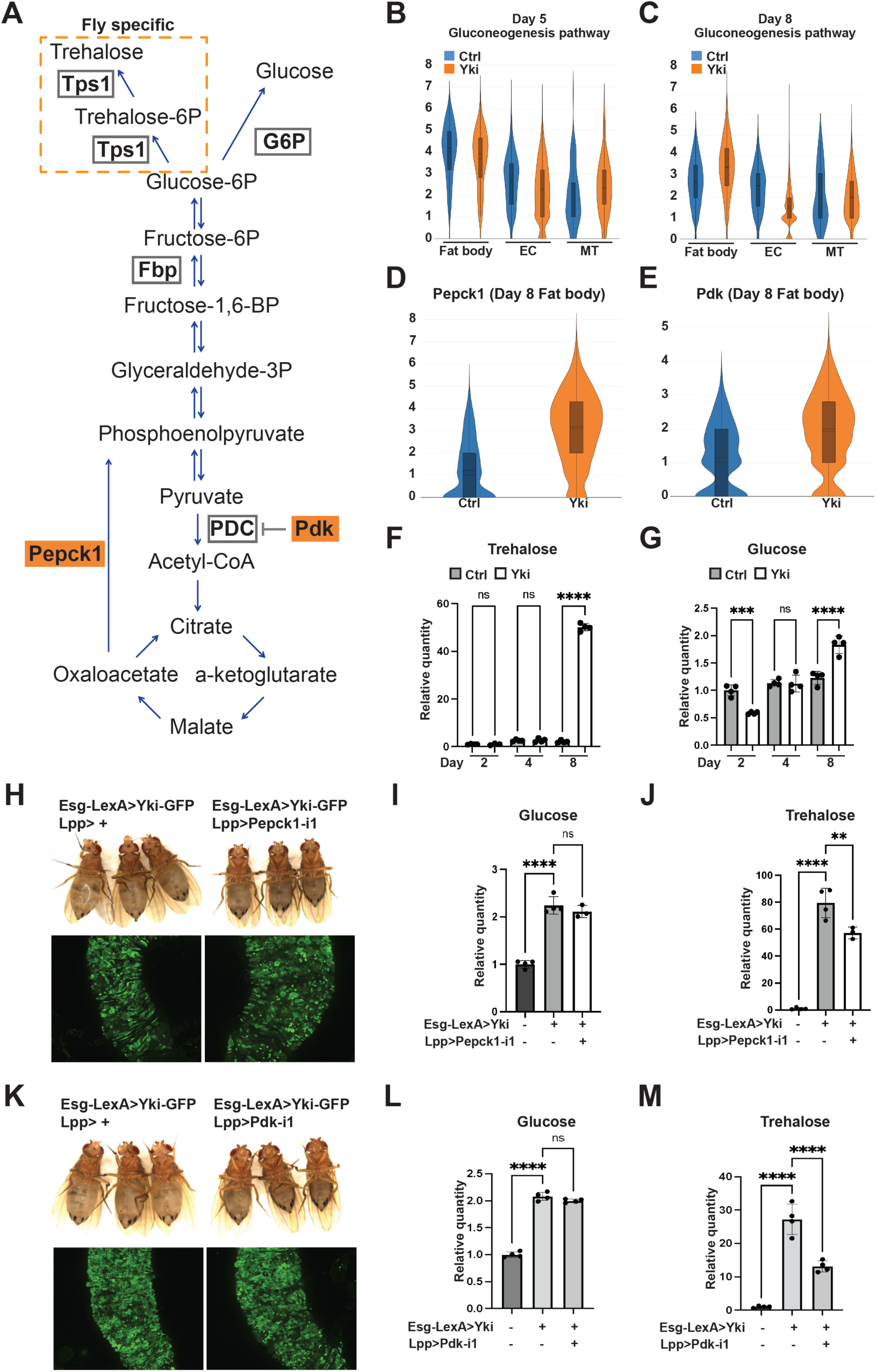
Increased expression of *Pepck1* and *Pdk* in the fat body of Yki flies stimulates gluconeogenesis. (A) Gluconeogenesis pathway in *Drosophila*. Expression levels of gluconeogenesis pathway genes at Day 5 **(B)** and Day 8 **(C)** in the fat body, EC, and malpighian tubule (MT) cell clusters visualized by violin plots. Expression levels of *Pepck1* **(D)** and *Pdk* **(E)** at Day 8 in fat body in control and Yki flies visualized by violin plots. Relative whole-body trehalose (**F**) and glucose (**G**) levels upon different tumor induction time. **(H)** Representative gut tumor and phenotype of Yki flies without and with fat body *Pepck1* depletion at Day 6. Relative whole-body glucose **(I)** and trehalose **(J)** levels of control flies, Yki flies, and Yki flies with fat body *Pepck1* depletion at Day 8. **(K)** Representative gut tumor and phenotype of Yki flies without and with fat body *Pdk* depletion at Day 6. Relative whole-body glucose **(L)** and trehalose **(M)** levels of control flies, Yki flies, and Yki flies with fat body *Pdk* depletion at Day 8. **p < 0.01, ***p < 0.001, ****p < 0.0001. Error bars indicate SDs. See also Figure S2.

### Elevated gluconeogenesis in Yki flies is independent of Akh and Insulin signaling

Insulin and glucagon are known to regulate gluconeogenesis in mammals (Exton, 1972). Likewise, insulin and glucagon-like adipokinetic hormone (Akh) control glucose metabolism in *Drosophila* (Chatterjee and Perrimon, 2021). Thus, we tested whether ImpL2 or Akh could regulate gluconeogenesis in the fat body of Yki flies. Depletion of the *Akh receptor* (*AkhR*) from the fat body of Yki flies had no significant effects on trehalose levels (Figure S3A), indicating that gluconeogenesis is regulated by another mechanism. For insulin signaling, inhibition of ImpL2 from Yki tumors reduces trehalose levels (Kwon et al., 2015). In addition, compared to Yki flies, removal of ImpL2 from Yki tumors, leads to an increase in insulin signaling in the fat body as indicated by a decrease in *InR* expression, a target gene of Insulin signaling that is up regulated when insulin signaling is low (Figure S3B) (Puig and Tjian, 2005). Interestingly, *Pepck1* expression levels were not reduced in the fat body of Yki flies with *ImpL2* depletion (Figure S3C), indicating that the abnormal level of gluconeogenesis in Yki flies is independent of insulin signaling.

### Systemic analysis of tumor-to-host organ communication

To search for a gut tumor secreted factor that regulates gluconeogenesis in the fat body, we analyzed tumor-host organ communication in Yki flies using FlyPhone, an integrated web-based resource for cell-cell communication predictions in *Drosophila* (Liu et al., 2022). First, we retrieved the list of genes encoding secreted proteins that were upregulated in the gut cell clusters following tumor induction. This included previously identified cachectic factors such as ImpL2, Upd3, and Pvf1 (Table S1) (Kwon et al., 2015; Song et al., 2019; Ding et al., 2021). Interestingly, although *yki*^act^ expression was induced solely in ISCs (Figure 3AB), the cachectic factors were also detected in ECs (Figure 3C-H, Table S1). Specifically, *ImpL2* and *Upd3* were upregulated in both ISCs and ECs (Figure 3C-F), while *Pvf1* was mainly expressed in ECs (Figure 3GH). This observation led us to analyze the inter-organ communication from not only tumor gut ISCs but also ECs to peripheral organs. FlyPhone generated a connectivity prediction graph of possible signaling interactions between gut tumor and host organs (Figure S3DE). By excluding pathways with weaker connectivity in fat body from Day 5 to Day 8, since the wasting phenotypes were more severe at Day 8, we identified a few candidates for regulators of gluconeogenesis in the fat body, including PVR RTK and JAK-STAT pathways (Figure 3IJ).

**Figure 3.**
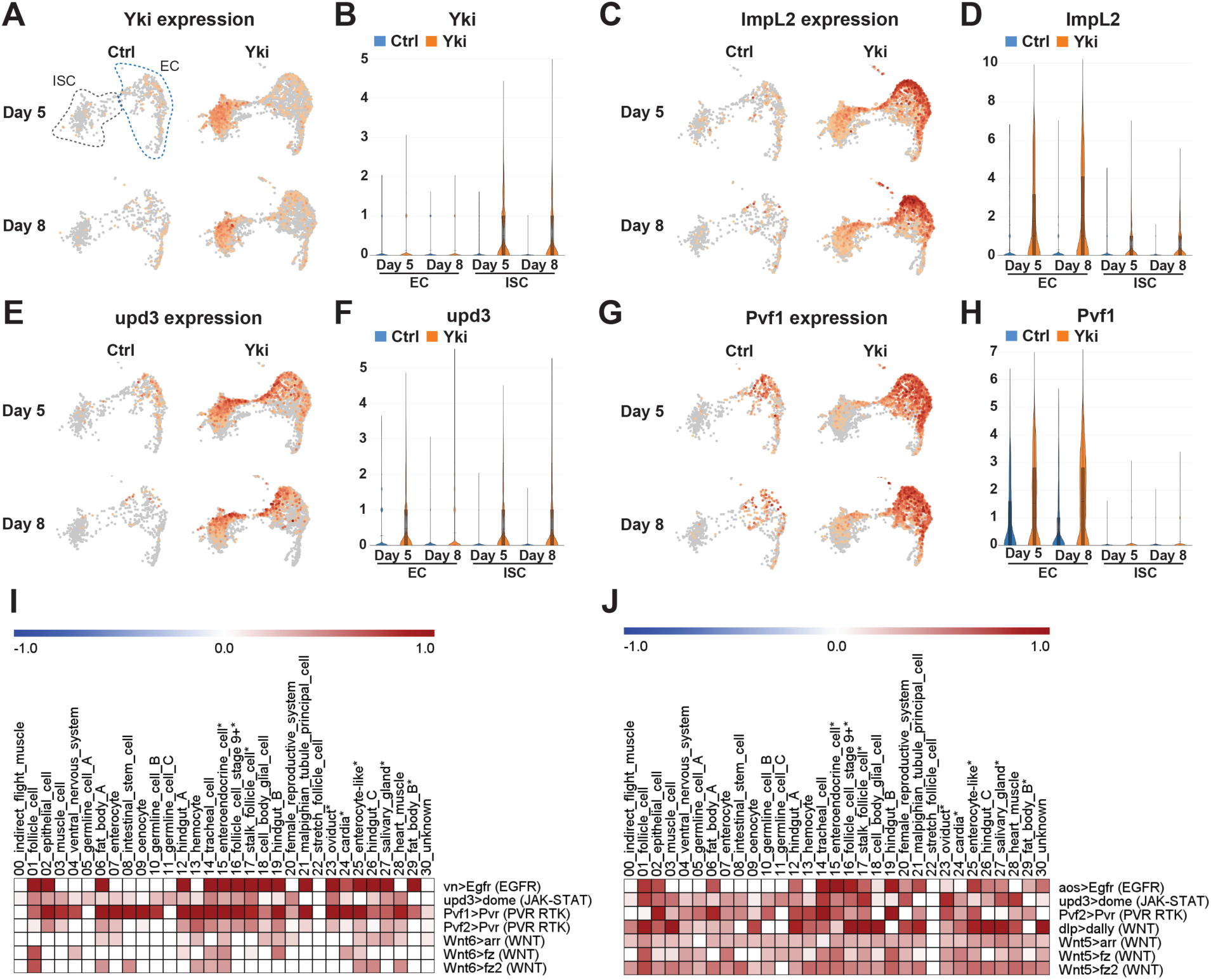
Analysis of perturbed signaling pathways in Yki flies. UMAP visualization showing enrichment of *Yki* **(A)**, *ImpL2* **(C)**, *Upd3* **(E)**, and *Pvf1* **(G)** expression in ISC and EC clusters. Expression levels of *Yki* **(B)**, *ImpL2* **(D)**, *Upd3* **(F)**, and *Pvf1* **(H)** in ISC and EC clusters represented by violin plots. Heatmap plots showing selected increased signaling from EC **(I)** and from ISC **(J)** upon tumor progression (Day 8 vs Day 5). Darkness reflects increased signaling. See also Figure S3.

### The JAK-STAT pathway stimulates hepatic gluconeogenesis

We first tested the PVR RTK signaling pathway because it has a role in metabolism regulation and has been shown to induce bloating in Yki flies (Barton, 2001; Song et al., 2019). However, activation of the Pvr pathway in wildtype flies through expression of *Pvf1* in ISC (*esg*>*Pvf1*) did not induce *Pepck1* or *Pdk* expression (Figure S4AB). Next, we examined the role of the JAK STAT pathway. Interestingly, overexpression of *Upd3* in wildtype fly ISCs (*esg>upd3)* increased fat body expression of *Pepck1* and *Pdk*, as well as *ImpL2* (Figure 4A-C). To further examine the regulation of *Pepck1* and *Pdk* by JAK-STAT, we activated JAK-STAT signaling specifically in the fat body through expression of a tagged constitutively active form of Stat92e (*Lpp*>*STAT act-HA*) (Figure S4C). Consistent with the observation of multiple STAT binding motifs in these regions, chromatin immunoprecipitation revealed that Stat92E physically associates with *Pepck1* and *Pdk* gene regions in the fat body (Figure 4D-F). These observations suggest that JAK-STAT signaling directly promotes fat body expression of *Pepck1* and *Pdk*, two genes essential for the elevated gluconeogenesis in Yki flies that contributes to the increased trehalose levels.

**Figure 4.**
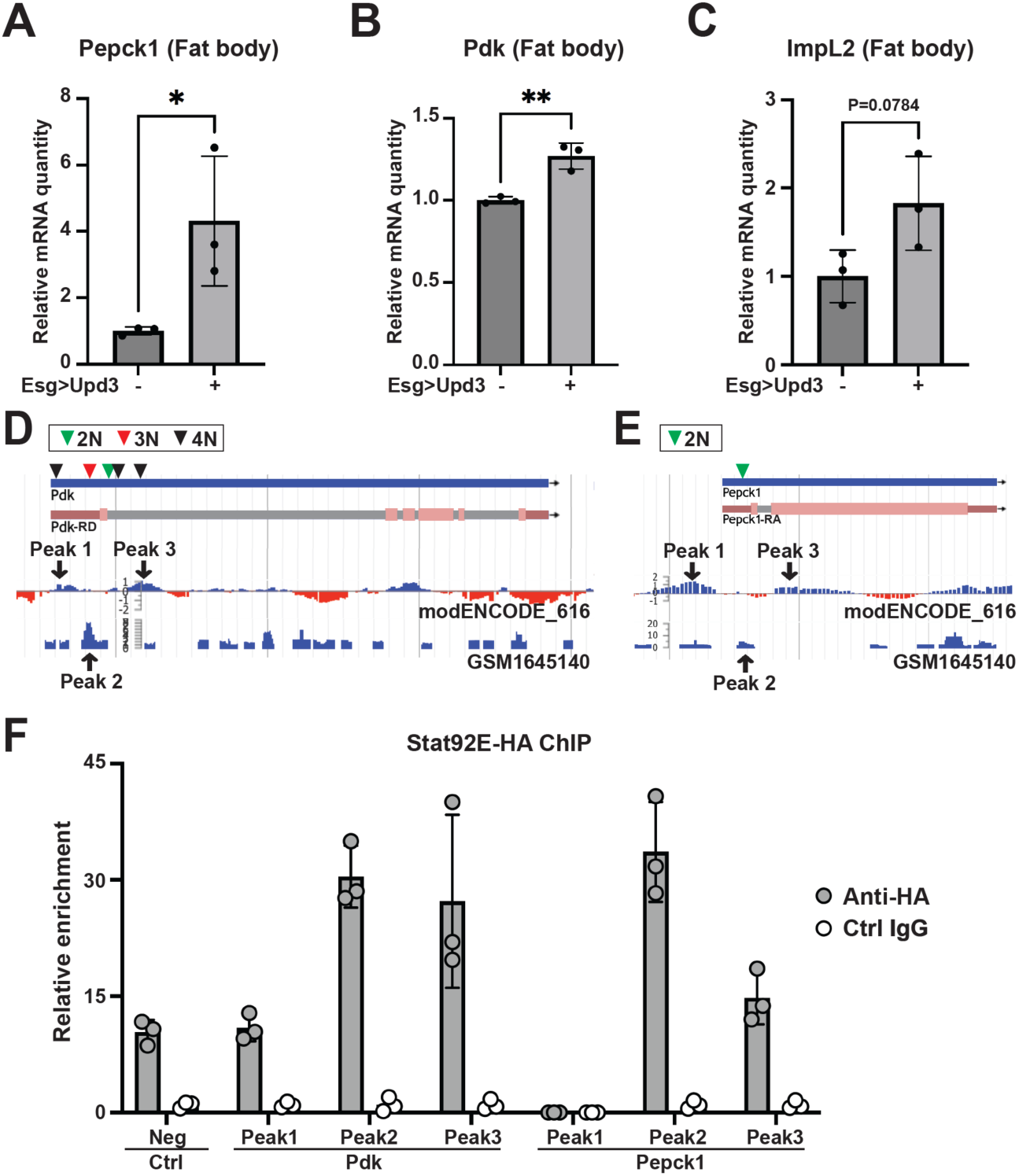
The Jak/Stat pathway regulates *Pepck1* and *Pdk* in the fat body. qRT-PCR analysis of *Pepck1* **(A),** *Pdk* **(B)**, and *ImpL2* **(C)** mRNA levels in fat body of flies without and with ISC *Upd3* expression at Day 8. Data retrieved from ChIP-seq database indicating enrichment of Stat92e binding at *Pdk* **(D)** and *Pepck1* **(E)** gene region, inverted triangle indicates STAT binding motif (2N: TTCNNGAA, 3N: TTCNNNGAA, 4N: TTCNNNNGAA). **(F)** Chromatin immunoprecipitation (ChIP) revealed the enrichment of HA-tagged Stat92E binding at *Pepck1* and *Pdk* gene region showed by fold changes related to control IgG at Day 8. A fragment of *Sam-S* with no Stat-binding sites was used as the negative control (Neg). *p < 0.05, **p < 0.01. Error bars indicate SDs. See also Figure S4.

### Conserved Jak/Stat pathway regulation of gluconeogenesis in mouse cancer cachexia models

To validate these findings in mammals, we utilized a well-established, inducible, genetically engineered mouse model of lung cancer (*Kras^LSL-G12D/+^*;*Lkb1^flox/flox^*, referred to as KL mice) (Ji et al., 2007; Goncalves et al., 2018). We induced tumors in KL mice through intranasal administration of adenovirus encoding for the Cre recombinase. 5–6 weeks after tumor induction, ∼60-70% of the mice develop cachexia as defined by a total body weight loss of more than 15% (Goncalves et al., 2018; Queiroz et al., 2022a). To decipher the cachexia-related alterations of glucose metabolism, we compared RNA-sequencing data from livers of KL mice with cancer anorexia-cachexia syndrome (CACS) or without it (NCACS) using gene set enrichment analysis (GSEA). We found that expression of gluconeogenesis genes is enriched in the livers of the cachectic mice (Figure S5AB). Interestingly, *Pck1* and *Pdk3*, the human homologs of fly *Pepck1* and *Pdk*, respectively, are among the genes upregulated in the livers of CACS KL mice (Figure 5A, Figure S5B). Upregulation of these genes in the liver leads to elevated hepatic production of glucose (Huang et al., 2002; Zhang et al., 2014; Yu et al., 2021), which is identical to what we observed in Yki flies. Importantly, liver expression levels of *Pck1* and *Pdk3* positively correlate with the weight loss of KL mice (Figure 5BC), suggesting a contribution of increased gluconeogenesis to the poor prognosis. Because the fly fat body is a liver-adipose hybrid organ, we also compared the transcriptome of white adipose tissue (WAT) between CACS and NCACS KL mice. Interestingly, we observed upregulation of different Pdks, *Pdk1* and *Pdk2*, in WAT in CACS mice (Figure 5D). Pdks inhibit the conversion of pyruvate to acetyl-CoA, which represses fatty acid synthesis from glucose, and thus might lead to higher blood glucose levels (Huang et al., 2002; Zhang et al., 2014). Interestingly, expression of *Pdk1* and *Pdk2* in WAT also positively correlates with the weight loss of KL mice (Figure 5EF), suggesting their contribution to cachexia. Furthermore, in both liver and WAT in CACS mice (Figure 5AD), we observed increased expression of *Igfbp3*, which is a functional equivalent of fly *ImpL2* and has been reported to antagonize insulin signaling in the context of cancer cachexia (Huang et al., 2016). Collectively, our data suggests that tumor induced liver-adipose tissue expression of *Pepck1*/*Pck1*, *Pdk*/*Pdk1-3*, and *ImpL2*/*Igfbp3* is conserved in Yki flies and KL mice.

**Figure 5.**
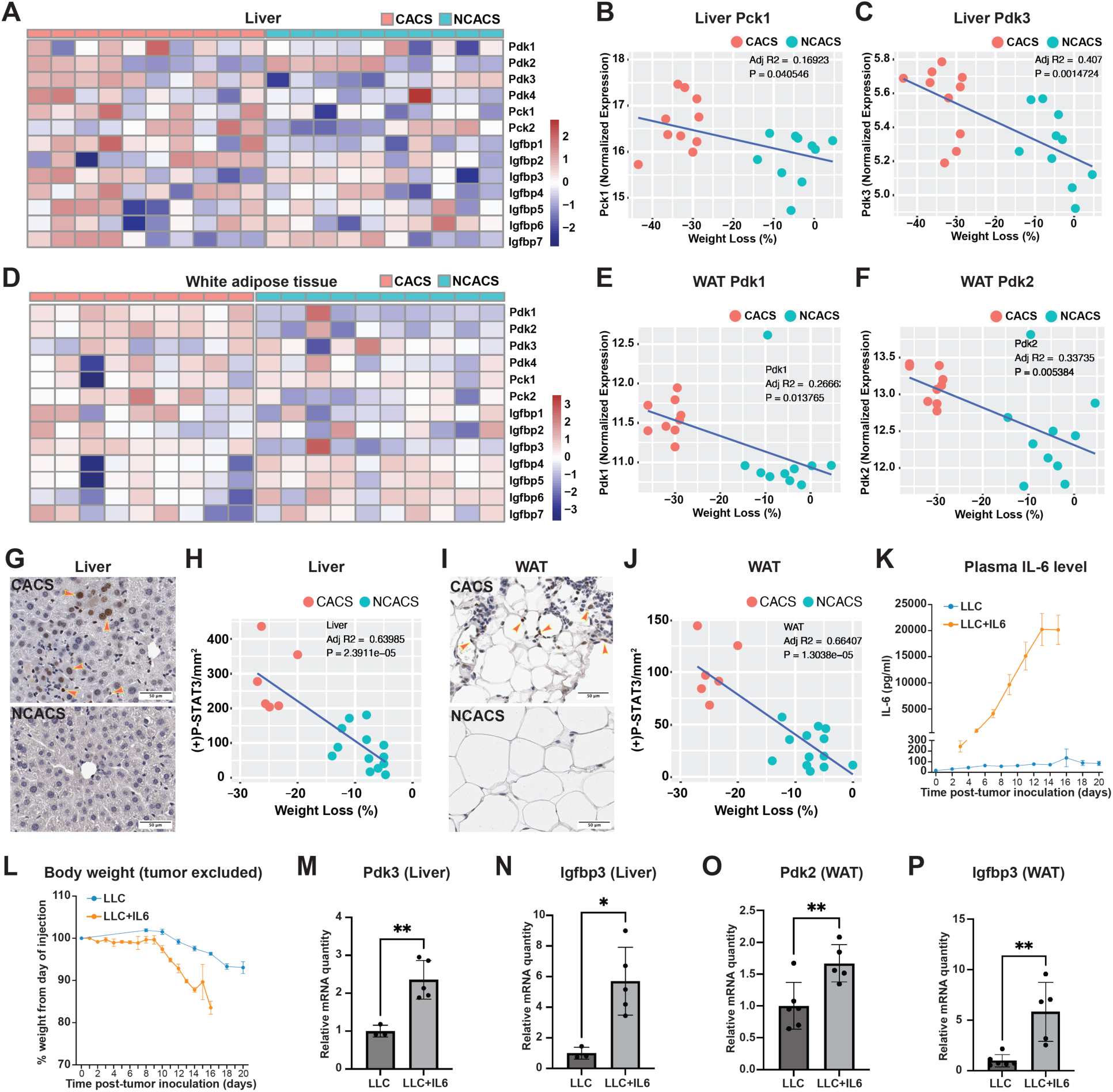
Conserved Jak/Stat pathway regulation of cachectic gene expression in hepatocytes and adipocytes. Heatmap plots showing expression levels of *Pdk1-4*, *Pck1-2*, and *Igfbp1-7* in liver **(A)** and WAT **(D)** of CACS and NCACS KL mice (Columns are showing individual animals). Correlation plots showing positive relations between liver *Pck1* expression **(B)**, liver *Pdk3* expression **(C)**, WAT *Pdk1* expression **(E)**, WAT *Pdk2* expression **(F)** and weight loss of KL mice. Immunohistochemistry (IHC) staining of p-STAT3 in liver **(G)** and WAT **(I)** of CACS and NCACS KL mice. Quantifications are shown in **(H)** and **(J)**, respectively. Plasma IL-6 levels **(K)** and body weight **(L)** of B6 mice injected with LLC cells without and with IL-6 expression. qRT-PCR analysis of liver *Pdk3* **(M)**, *Igfbp3* **(N)**, and WAT *Pdk2* **(O)**, *Igfbp3* **(P)** mRNA levels of B6 mice injected with LLC cells without and with IL-6 expression. *p < 0.05, **p < 0.01. Error bars indicate SDs. See also Figure S5.

Next, we hypothesized that the cachexia-related expression of *Pck1*, *Pdk1-3*, and *Igfbp3* in CACS KL mice is regulated through the same mechanism of IL-6/JAK-STAT signaling. Indeed, IL-6 is only detectable in the serum of CACS but not NCACS KL mice (Goncalves et al., 2018). Supporting this hypothesis, we observed higher levels of phospho-STAT3 (p-STAT3) in liver and WAT in CACS KL mice (Figure 5G-J). To examine the regulation of these genes by IL-6, we compared B6 mice injected with Lewis Lung Carcinoma (LLC) cells engineered to express or not express IL-6 (Figure 5K). Notably, mice injected with LLC cells producing IL-6 (LLC+IL6) displayed rapid weight loss (Figure 5L). Consistent with our observations in KL mice, liver expression of *Pdk3* and *Igfbp3*, and WAT expression of *Pdk2* and *Igfbp3*, were upregulated in mice injected with LLC cells producing IL-6 (Figure 5M-P). We did not observe an upregulation of *Pck1* in liver in LLC+IL6 mice (Figure S5C), suggesting that *Pck1* induction may require additional conditions specific to KL tumors. As PDKs promote gluconeogenesis and represses glycolysis and lipogenesis in a cell-autonomous manner (Huang et al., 2002; Tao et al., 2013; Zhang et al., 2014), our data suggest that JAK-STAT signaling induces cancer cachexia-related hyperglycemia through targeting of multiple organs.

### Hepatic inhibition of JAK-STAT signaling rescues the cachectic symptoms of Yki flies

Given that cancer cachexia is a major factor that affects the health status and survival of patients (Tisdale, 2009; Liu et al., 2022), we wanted to test whether inhibition of hepatic JAK STAT signaling in Yki flies affect cachectic symptoms. Strikingly, blocking JAK-STAT signaling in Yki fly fat bodies through depletion of *hop/JAK* or *Stat92e* reduced *Pepck1*, *Pdk*, and *ImpL2* expression levels with no obvious effects on the gut tumor cells (Figure 6A-C, Figure S6AB). Further, depletion of *hop/JAK* and *Stat92e* inhibited the bloating and elevated whole-body trehalose levels in Yki flies, but had no effects on glucose, glycogen, or TAG levels (Figure 6DE, Figure S6CD), as observed following the knockdown of either *Pepck1* or *Pdk* in the fat body in Yki flies (Figure 3, Figure S3). Thus, inhibition of JAK-STAT signaling in the fat body is sufficient to repress expression of these cachectic genes and to attenuate the cachexia phenotypes. Finally, we characterized how elevated JAK-STAT signaling and hepatic gluconeogenesis in Yki flies affect the overall mobility and viability of Yki flies. Inhibition of JAK-STAT signaling (*hop/JAK* depletion) and gluconeogenesis (*Pdk* depletion) in Yki fly fat bodies both restored climbing ability and improved overall survival of Yki flies (Figure 6FG), suggesting that hepatic gluconeogenesis is the major cause of the JAK-STAT signaling induced cachectic symptoms of Yki flies.

**Figure 6.**
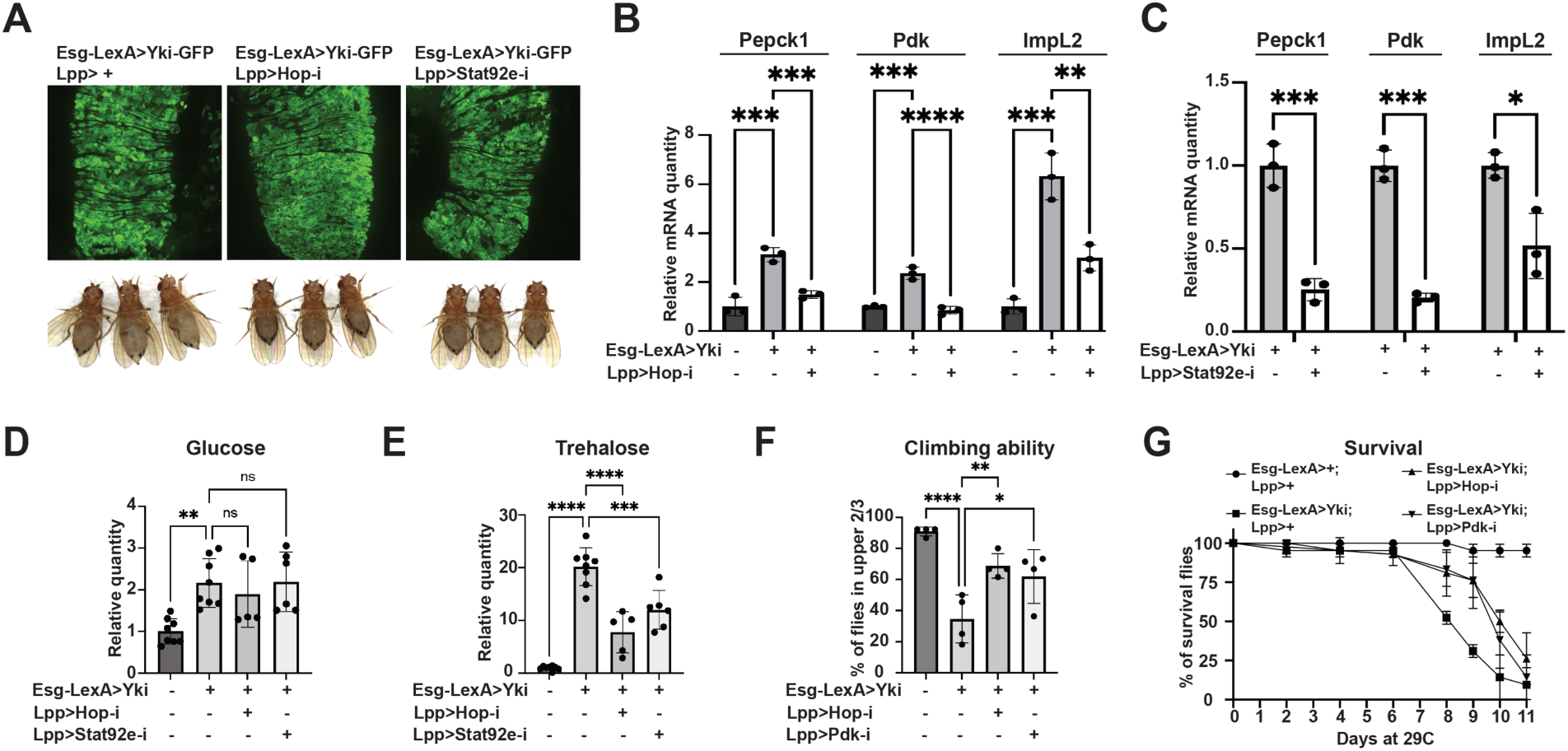
Cachectic role of tumor-induced JAK-STAT signaling. **(A)** Representative gut tumor and phenotypes of Yki flies without and with fat body *hop* or *Stat92e* depletion at Day 6. **(B)** qRT-PCR analysis of *Pepck1*, *Pdk*, and *ImpL2* mRNA levels in the fat body of control flies, Yki flies, and Yki flies with fat body *hop* depletion at Day 8. **(C)** qRT-PCR analysis of *Pepck1*, *Pdk*, and *ImpL2* mRNA levels in the fat body of Yki flies without or with fat body *Stat92e* depletion at Day 8. Relative whole-body glucose **(D)** and trehalose **(E)** levels of control flies, Yki flies, and Yki flies with fat body *hop* or *Stat92e* depletion at Day 6. Climbing ability at Day 6 **(F)** and survival curve **(G)** of control flies, Yki flies, and Yki flies with fat body *hop* or *Pdk* depletion. *p < 0.05, **p < 0.01, ***p < 0.001, ****p < 0.0001. Error bars indicate SDs. See also Figure S6.

## Discussion

Hyperglycemia is the earliest metabolic abnormality observed in cancer patients and it is generally agreed that the abnormal blood glucose levels are caused by cancer-induced insulin resistance (Rohdenburg et al., 1919; Tayek, 1992; Honors and Kinzig, 2012). In this study, we leveraged single nuclei transcriptomics to systemically investigate tumor induced host metabolism reprogramming and tumor-host organ communication in a *Drosophila* cancer cachexia model, and discovered a previously unknown but conserved cachectic role of Upd3/IL 6 in induction of hepatic gluconeogenesis. Our findings add to the current knowledge of the pathological basis of IL-6 in cancer cachexia.

### Full body snRNASeq analysis provides a comprehensive understanding of *yki*^act^ tumor induced cachectic factors

Previous studies have shown that Yki flies have increased expression of cachectic factors including *ImpL2, Upd3*, and *Pvf1* (Kwon et al., 2015; Song et al., 2019; Ding et al., 2021). Because the expression levels of these factors were detected through bulk RNAseq and qPCR, it was not clear whether they were derived from ISCs or other gut cell types. Our snRNAseq data indicates that *ImpL2* and *Pvf1* are mainly induced in ECs in the tumorous gut (Table S1, Figure 3CD, GH). In addition, we found increased *Upd3* expression in both ISCs and ECs in tumorous guts (Figure 3EF). Notably, a subset of cells closes to the junction of ISC and EC clusters displayed high levels of *Upd3* expression (Figure 3E). Finally, another cachectic factor specifically expressed in *yki*^act^ tumor gut ECs is *matrix metalloproteinase 2* (*Mmp2*) (Table S1). *Mmp2* was previously identified in a *Drosophila* larval tumor model and shown to induce muscle wasting (Lodge et al., 2021); however, whether Mmp2 contributes to muscle degeneration in Yki flies remains to be tested. Altogether, these data highlight that the cellular heterogeneity of tumors may drive the production of different cachectic factors.

### Decreased anabolism and elevated gluconeogenesis contribute to cachexia in Yki flies

Body mass is controlled by homeostasis between catabolism and anabolism (McCarthy and Esser, 2010). Our study indicates that decreased anabolism and elevated gluconeogenesis underly the loss of body mass in Yki flies. First, body mass loss of Yki flies appears to originate from decreased anabolism rather than elevated catabolism, as we observed decreased Lipid and glycogen production in Yki flies at Day 8, a time point where degradation processes were not enhanced (Figure 1, Figure S1). The systemic decrease in glycolysis in Yki flies may further inhibit anabolism through reducing fuel availability (Figure 1, Figure S1). Second, the sole inhibition of hepatic gluconeogenesis in Yki flies increased their climbing abilities and survival rates (Figure 6FG), indicating its contribution to the severity of the cachexia phenotype. Notably, gluconeogenesis is an energy consuming process that increases body energy expenditure (Blackman, 1982). Consistent with our data, increased energy expenditure is another accelerator of body wasting reported in cancer patients (Hyltander et al., 1991; Cao et al., 2010; Friesen et al., 2015). Although tumors require energy for rapid cell proliferation, it is unlikely that the metabolic demands of tumors play a major role in generating negative energy balance since the energy expenditure of tumors are often minor (<5%) when cachexia occurs (Keller, 1993; Cairns et al., 2011; Fearon et al., 2012). Therefore, tumor-induced energy expenditure in peripheral organs, such as excessive hepatic gluconeogenesis, may be an important stimulator of cachexia (Stein, 1978; Blackman, 1982; Holroyde et al., 1984; Keller, 1993; Bongaerts et al., 2006).

### Upd3/IL-6 controls hepatic glucose production independent of insulin and glucagon signaling

IL-6 has been commonly viewed as a proinflammatory cytokine; however, growing evidence suggests a broader role for IL-6 in regulating glucose homeostasis (Lehrskov and Christensen, 2019). For instance, an early study indicated that incubating rat hepatocyte primary cultures with IL-6 can induce gluconeogenesis (Blumberg et al., 1995). In particular, IL 6 neutralization attenuates elevated hepatic glucose production in high-fat-fed rats and IL-6 infusion promotes hepatic glucose production in control mice (Perry et al., 2015). A recent study suggested that stress induced IL-6 is required for an acute hyperglycemia for adaptive “fight or flight” responses (Qing et al., 2020). In our study, tumor-induced Upd3/IL-6 promotes gluconeogenesis gene expression through activation of hepatic JAK-STAT signaling in both mouse and fly. In animal and humans, gluconeogenesis is tightly controlled by insulin and glucagon (Exton, 1972; Kraus-Friedmann, 1984). Interestingly, restoration of insulin signaling and inhibition of Akh (glucagon-like hormone) signaling in Yki flies showed no effect on the increased gluconeogenesis (Figure S3A-C), suggesting that JAK-STAT signaling overrides the normal physiological regulation of gluconeogenesis. Notably, IL-6 induced hepatic *Pck1* expression seems to be context dependent, as evidenced from increased expression of *Pck1* in STAT3 knock out mice and STAT3 -dependent inhibition of *Pck1* expression in HepG2 cells (Inoue et al., 2004; Nie et al., 2009; Ramadoss et al., 2009). However, STAT3 does bind to the promoter region of *Pck1* (Ramadoss et al., 2009). These observations suggest that additional regulators, which may be present in condition of certain cancers, are involved in controlling *Pck1* expression together with IL-6. Nevertheless, Upd3/IL-6/JAK-STAT dependent induction of *Pdk*/*Pdks* expression is consistent among all the conditions we tested, including both fly and mouse samples. Altogether, these observations indicate a currently underestimated role of Upd3/IL-6 in regulating hepatic gluconeogenesis.

### Novel cachectic role of PDK3 in cancer

Humans and rodents have four PDKs, *Pdk1-4* (Popov et al., 1997; Bowker-Kinley et al., 1998), which have different expression pattern across tissues (Bowker-Kinley et al., 1998). In the rat liver, Pdk2 is abundant and Pdk4 is expressed at a much lower level (Bowker-Kinley et al., 1998). In mice, *Pdk1* and *Pdk2* are both highly expressed in the liver (Klyuyeva et al., 2019). Among them, *Pdk2* and *Pdk4* show increased expression in response to starvation or diabetes *Pdk2* is upregulated in liver and kidney, whereas *Pdk4* is highly induced in heart, muscle, kidney, and slightly in liver (Wu et al., 1998, 2000; Sugden et al., 2000). Notably, *Pdk3* expression was not detectable in the liver in humans and rodents, neither in normal conditions nor in starvation and diabetes (Gudi et al., 1995; Bowker-Kinley et al., 1998; Klyuyeva et al., 2019), indicating a distinct regulatory mechanism and physiological role of *Pdk3*. In support of this, PDK3 displays many unusual biochemical characteristics: 1) Among the recombinant PDK isoenzymes, PDK3 exhibits the highest catalytic activity, 25-fold higher than the activity of PDK2 (Bowker-Kinley et al., 1998); 2) Activation of PDK3 does not depend on the levels of NADH and acetyl-CoA, which are required for activation of other PDKs (Bowker-Kinley et al., 1998); and 3) PDK3 is the PDK least sensitive to the inhibition of pyruvate, 40-fold less sensitive than PDK2 for this feed-back inhibition (Bowker Kinley et al., 1998; Baker et al., 2000). As such, in our mouse cancer cachexia models, IL 6/JAK-STAT signaling induced the expression of a highly efficient, autonomously activated, and almost non-repressible PDK in the liver, which may facilitate an intensive and prolonged inhibition of acetyl-CoA production from pyruvate, leading to insulin resistance consequences including increased gluconeogenesis, reduced energy release from the tricarboxylic acid (TCA) cycle, and decreased fatty acid synthesis. These observations suggest that the IL-6/JAK-STAT signaling-dependent induction of hepatic expression of *Pdk3* is a previous unknown mechanism of cancer cachexia.

### Upd3/IL-6 induces insulin resistance through multiple mechanisms

We reported previously that fly tumor secreted Upd3 induces insulin resistance through induction of *ImpL2* expression in peripheral tissues (Ding et al., 2021). In this study, we demonstrate that Upd3 targets the fat body, a liver-adipose hybrid organ, to induce insulin resistance through additional mechanisms. Upd3/JAK-STAT signaling induced fat body expression of *Pepck1* and *Pdk* promotes hepatic gluconeogenesis which mimic hepatic insulin resistance in mammals (Bock et al., 2007; Meshkani and Adeli, 2009; Petersen and Shulman, 2018). Besides facilitating gluconeogenesis, *Pdk* represses glycolysis and lipogenesis in a cell-autonomous manner, which contributes to local insulin insensitivity (Huang et al., 2002; Zhang et al., 2014). Importantly, we observed identical gene expression regulations in mouse models, as IL-6 upregulates *Igfbp3* in both liver and WAT, *Pck1* and *Pdk3* in liver, and *Pdk1-2* in WAT. Collectively, Upd3/IL-6 targets multiple peripheral tissues to stimulate local and systemic insulin resistance which contribute to the metabolic dysregulation observed in cachexia.

In conclusion, we provide new insights into the pathogenesis of cancer cachexia leveraging a multi-model approach. We systematically deciphered the metabolic dysregulation associated with cancer cachexia through body-wide single-cell transcriptome profiling in flies, which identified the cachectic role of gluconeogenesis. We further supported this finding with results of preclinical mouse models. This approach facilitates our uncovering of the conserved pathogenic role of Upd3/IL-6/JAK-STAT signaling in cancer-associated insulin resistance, providing a potential new therapeutic avenue of targeting hepatic gluconeogenesis in IL-6 related cancer cachexia.

## Acknowledgments

We thank Rich Binari, Pedro Saavedra, Joshua Li, Patrick Jouandin, Ismail Ajjawi, and all members of the Perrimon Lab for their critical suggestions and help on this research. We thank Stephanie Mohr for comments on the manuscript. We thank Hongjie Li and Sudhir Gopal Tattikota for advice on single nuclei sequencing, Paula Montero Llopis and Microscopy Resources on the North Quad (MicRoN) core facility at Harvard Medical School for advice and help on confocal imaging, Jodene Moore and the Systems Biology FACS core facility at Harvard Medical School for advice and help on flow cytometry, Biopolymers facility at Harvard Medical School for sequencing, and the *Drosophila* RNAi Screening Center (DRSC) and Bloomington *Drosophila* Stock Center (BDSC) for providing fly stocks used in this study. We thank Erika Bach for the generous gift of Stat92E fly stocks.

This article is subject to HHMI’s Open Access to Publications policy. HHMI lab heads have previously granted a nonexclusive CC BY 4.0 license to the public and a sublicensable license to HHMI in their research articles. Pursuant to those licenses, the author-accepted manuscript of this article can be made freely available under a CC BY 4.0 license immediately upon publication.

## Materials and Methods

### Drosophila strains

All flies were kept on standard cornmeal fly food supplemented with yeast and agar. Crosses were grown at 18°C to inactivate Gal4 and LexA. Adult offspring flies were collected within 48 hours after emerging, kept at 18°C for another 24 hours and then incubated at 29°C for indicated days to induce transgene expression (e.g., “Day 8” indicates flies were collected after 8 days of transgene expression induction). Flies were flipped onto fresh food every 2 days. Stocks used in this study include *esg-Gal4, tub-Gal80ts, UAS-GFP* (Kwon et al., 2015), *esg-LexA::GAD* (BDSC 66632), *tub-Gal80ts, Lpp-Gal4* (Song et al., 2017), *CG31272-Gal4* (BDSC 76171), *UAS-Yki3SA* (Oh and Irvine, 2009), *LexAop-Yki3SA-GFP* (Saavedra et al., 2021), *UAS-Pepck1-RNAi* (RNAi-1 BDSC 65087, RNAi-2 VDRC 50253), *UAS-Pdk-RNAi* (RNAi-1 BDSC 35142 & RNAi-2 28635), *UAS-Hop-RNAi* (BDSC 32966), *UAS-Stat92e-RNAi* (BDSC 33637), *UAS-ImpL2-RNAi* (NIG 15009R3), *UAS-AkhR-RNAi* (BDSC 51710), *UAS-HA-Stat92E* dominant-active form (Ekas et al., 2010), *UAS-Upd3* (Woodcock et al., 2015), and *UAS-Pvf1*(Xu et al., 2022). *w1118* was used as control. Female flies are used in all experiments as they showed more significant and consistent bloating phenotype.

## Genotypes used in this study

| Figure |  |
| --- | --- |
| 1&S1<br>2B-G<br>S2A-F | *esg-GAL4, tub-GAL80TS > UAS-GFP*<br>*esg-GAL4, tub-GAL80TS > UAS-GFP, UAS-yki<sup>3SA</sup>* |
| 2H-J<br>S2G-L | *esg-LexA, tub-GAL80TS > +; Lpp>+*<br>*esg- LexA, tub-GAL80TS > LexAop-yki<sup>3SA</sup>-GFP; Lpp>+*<br>*esg- LexA, tub-GAL80TS > LexAop-yki<sup>3SA</sup>-GFP; Lpp>Pepck-i* |
| 2K-M<br>S2M-Q | *esg-LexA, tub-GAL80TS > +; Lpp>+*<br>*esg- LexA, tub-GAL80TS > LexAop-yki<sup>3SA</sup>-GFP; Lpp>+*<br>*esg- LexA, tub-GAL80TS > LexAop-yki<sup>3SA</sup>-GFP; Lpp>Pdk-i* |
| 3A-J<br>S3DE | *esg-GAL4, tub-GAL80TS > UAS-GFP*<br>*esg-GAL4, tub-GAL80TS > UAS-GFP, UAS-yki<sup>3SA</sup>* |
| S3A | *esg-LexA, tub-GAL80TS > +; Lpp>+*<br>*esg-LexA, tub-GAL80TS > LexAop-yki<sup>3SA</sup>-GFP; Lpp>+*<br>*esg-LexA, tub-GAL80TS > LexAop-yki<sup>3SA</sup>-GFP; Lpp>AkhR-i* |
| S3BC | *esg-GAL4, tub-GAL80TS > UAS-GFP, UAS-yki<sup>3SA</sup>*<br>*esg-GAL4, tub-GAL80TS > UAS-GFP, UAS-yki<sup>3SA</sup>, UAS-Impl2-i* |
| 4A-C | *esg-GAL4, tub-GAL80TS > UAS-GFP*<br>*esg-GAL4, tub-GAL80TS > UAS-GFP, UAS-Upd3* |
| S4AB | *esg-GAL4, tub-GAL80TS > UAS-GFP*<br>*esg-GAL4, tub-GAL80TS > UAS-GFP, UAS-Pvf1* |
| 4F<br>S4C | *Lpp-GAL4, tub-GAL80TS > +*<br>*Lpp-GAL4, tub-GAL80TS > UAS-STAT-act* |
| 5A-J<br>S5AB | *Kras<sup>LSL-G12D/+</sup>; Lkb1<sup>flox/flox</sup>* |
| 5K-P<br>S5C | *C57BL/6J* |
| 6A-E<br>S6 | *esg-LexA, tub-GAL80TS > +; Lpp>+*<br>*esg-LexA, tub-GAL80TS > LexAop-yki<sup>3SA</sup>-GFP; Lpp>+*<br>*esg-LexA, tub-GAL80TS > LexAop-yki<sup>3SA</sup>-GFP; Lpp>Hop-i*<br>*esg-LexA, tub-GAL80TS > LexAop-yki<sup>3SA</sup>-GFP; Lpp>Stat92e-i* |
| 6FG | *esg-LexA, tub-GAL80TS > +; Lpp>+*<br>*esg-LexA, tub-GAL80TS > LexAop-yki<sup>3SA</sup>-GFP; Lpp>+*<br>*esg-LexA, tub-GAL80TS > LexAop-yki<sup>3SA</sup>-GFP; Lpp>Hop-i*<br>*esg-LexA, tub-GAL80TS > LexAop-yki<sup>3SA</sup>-GFP; Lpp>Pdk-i* |

## Whole-body single nuclei profiling of adult flies

Single nuclei suspensions were prepared as described by Fly Cell Atlas (Li et al., 2022). Whole body flies without heads were flash-frozen using liquid nitrogen and were homogenized in 1ml dounce in buffer of 250 mM sucrose, 10 mM Tris pH 8.0, 25 mM KCl, 5mM MgCl, 0.1% Triton-X, 0.5% RNasin plus (Promega, N2615), 1X protease inhibitor (Promega, G652A), 0.1 mM DTT, then filtered through 40 um cell strainer and 40 um Flowmi (BelArt, H13680-0040). Samples were centrifuged, washed, and resuspended in 1 X PBS with 0.5% BSA and 0.5% RNasin plus. The suspension was filtered again with 40 um Flowmi immediately before FACS sorting. Nuclei were stained with DRAQ7TM Dye (Invitrogen, D15106) and sorted using Sony SH800Z Cell Sorter at Systems Biology Flow Cytometry Facility at Harvard Medical School. After sorting, nuclei were collected and re-suspend at 700-800 cells/µl in 1 X PBS buffer with 0.5% BSA and 0.5% RNasin plus.

snRNAseq was performed according to the 10X genomics protocol (Chromium Next GEM Single Cell 3’_v3.1_Rev_D). Briefly, 16,000 nuclei were loaded on Chip G for each reaction. 2 reactions of control flies and 3 reactions of Yki tumor flies were processed for each time point (Day 5 and Day 8). Sequencing was conducted using Illumina NovaSeq 6000 S1 at Harvard Medical School Biopolymers Facility and reads were aligned to *Drosophila melanogaster* BDGP6.32. We processed the snRNAseq data using Cellranger count pipeline 6.1.1 and generated the feature-barcode matrices. The matrices from different samples were normalized by equalizing the read depth and aggregated into a single feature-barcode matrix using Cellranger aggr pipeline. In total, 122,898 cells were profiled including 25,146 control fly cells and 42,375 tumor fly cells at Day 5, and 19,050 control fly cells and 36,327 tumor fly cells at Day 8. We visualized the cell clusters and gene expression levels using Loupe Browser 6.

## Mouse models

The *Kras^G12D/+^*; *Lkb1^f/f^* mice have been described before (Ji et al., 2007), and were further backcrossed to FVB mice. Tumor induction in adult FVB mice (12 to 20-week-old) was achieved by intranasal administration of 75 μL of PBS containing 2.5 × 10^7^ pfu of Adenovirus CMV-Cre (Ad5CMV-Cre) obtained from the University of Iowa Gene Transfer Vector Core (Iowa City, IA) and 1 mM CaCl2. We had previously defined mice as CACS if they lost more than 15% of body weight from their peak weight over the course of the experiment, otherwise they were classified as NCACS (Goncalves et al., 2018; Queiroz et al., 2022). C57BL/6J mice were obtained from the Jackson Laboratory (Strain #000664). After a week of acclimation, 2 x 10^6^ Lewis Lung Carcinoma (LLC) cells or LLC cells edited to produce IL-6 (LLC-IL6) were subcutaneously inoculated into their right flank. Mice were kept in pathogen-free conditions on a 24 hour 12:12 light-dark cycle. All animal experiments were approved by the Institutional Animal Care and Use Committee (IACUC) at Cold Spring Harbor Laboratory (CSHL) and were conducted in accordance with the National Institutes of Health Guide for the Care and Use of Laboratory Animals. Body weights and clinical signs of cachexia were monitored on a daily basis. Handling was kept to a minimum. Mice were sacrificed when tumor size exceeded 2 cm length, when weight loss exceeded 15% from peak weight, or when showing clinical signs of discomfort indicative of cachectic endpoint as stated by the Animal Cachexia Score (ACASCO): piloerection, diarrhea or constipation, hunched posture, tremors, and closed eyes. Death was confirmed by cervical dislocation. Mice injected with LLC-IL6 cells reached >15% bodyweight loss endpoint. Mice in the LLC group were sacrificed 22-24 days after injection of the LLC cell line, when tumors reached 2 cm in length. LLC group mice did not reach cachectic endpoint but did exhibit a mild cachectic phenotype, characterized by reduced adipose and muscle tissue mass, and splenomegaly compared to non-tumor bearing control group mice.

## Plasma measurements from C57BL/6J mice

Tail vein bleeds were performed using a scalpel via tail venesection without restraint. Plasma samples were collected using heparin-coated hematocrit capillary tubes to avoid coagulation and were processed as follows: centrifuge spin at 14,000 rpm for 5 min at 4°C, snap frozen in liquid nitrogen and stored at -80°C. IL-6 levels were measured from plasma using the mouse IL 6 Quantikine ELISA Kit (#M6000B; R&D Systems).

## Lewis Lung Carcinoma (LLC) cell line

LLC cells were cultured in complete growth medium consisting of Dulbecco’s Modified Eagle Medium (DMEM) (#10027CV; Corning) containing 10% of Heat-Inactivated Fetal Bovine Serum (FBS) (#10-438-026; Thermo Fisher) and 1x Penicillin-Streptomycin solution (#15-140-122; Thermo Fisher) under sterile conditions. 1x Trypsin-EDTA (#15400054; Thermo Fisher) was used for cell dissociation. Cells were resuspended in FBS-free DMEM and viable cells were counted using a Vi-Cell counter prior to subcutaneous injection of 2x10^6^ viable cells diluted in 100μL DMEM into the right flank of each C57BL/6J mouse.

LLC cells were edited to constitutively produce interleukin-6 (IL-6). LLC cells were seeded into 24-well plates with 50,000 cells per well. After 24 hours, they were transfected with a total of 500ng of plasmid (comprising 2.5:1 PB-IL6 plasmid and PBase plasmids) using Lipofectamine 3000 (Thermo Fisher) according to the manufacturer’s protocol. PBase plasmid was obtained from System Biosciences (#PB210PA-1) and PB-IL6 was obtained from VectorBuilding comprising mouse Il6 cDNA driven by EF1a promoter and flanked by piggyBac elements. After 48 hours, the media was changed and replaced with DMEM media supplemented with 3 μg/ml puromycin. After 14 days of antibiotic selection, the media was replaced with DMEM media for 24 hours, followed by isolation of monoclonal populations by serial dilutions in a 96-well plate.

To identify clones with constitutive IL-6 expression, we measured IL-6 in the cell supernatant for each clone using the Mouse IL-6 ELISA Kit (#ab222503; Abcam).

## Bulk RNA-Sequencing from the KL livers

Total RNA was extracted from the liver using TRIzol (Thermo Fisher), followed by a clean-up step using RNeasy kit (Qiagen). One microgram of total RNA from each sample was submitted to the WCM Genomics Resources Core Facility. Raw sequenced reads were aligned to the mouse reference GRCm38 using STAR (v2.4.1d, 2-pass mode) aligner, and raw counts were obtained using HTSeq (v0.6.1). Differential expression analysis, batch correction and principal component analysis (PCA) were performed using R Studio Version 4.2.2 and DESeq2 (v.1.38.3). Gene set enrichment analysis (GSEA) analysis was performed with the R package fGSEA (<u>10.18129/B9.bioc.fgsea</u>), using the Reactome pathway database contained in the 2022 release of Mouse Molecular Signatures Database from the Broad Institute (https://www.gsea-msigdb.org/gsea/msigdb/mouse/collections.jsp).

## Quantitative RT-PCR

For fly samples, Nucleospin RNA kit (Macherey-Nagel) was used to extract RNA. cDNA was synthesized using iScript™ cDNA Synthesis Kit (Bio-Rad, 1708890) according to the manufacturer’s protocol. qPCR was performed with Thermal Cycler CFX 96 Real-Time System qPCR machine using iQ™ SYBR® Green Supermix (Bio-Rad). RP49 and CG13220 were used as housekeeping gene. Primers used for qPCRs are Hop (CAATTCCGTTGCACTCGGCG & GGCTCCAGGGAATCGTGTGG) Stat92e (CATCCTTTATTGGCTTCCAATGCTG & GCAAACTCTGCCCTGATGACTC) InR (GAAATGGCCACCTTAGCGGC & GGACAATTTTCCGGCCTCTCC) ImpL2 (AAGAGCCGTGGACCTGGTA & TTGGTGAACTTGAGCCAGTCG) Pdk (GGCAAGGAGGACATCTGTGT & AATGTCGCCATGGAAATAGC) Pdk 2^nd^ (GTTGCCGCTCTTCGTCGCAC & CTCTTCGCAGGACGTTGACC) Pepck1 (CGCCCAGCGACATGGATGCT & GTACATGGTGCGACCCTTCA) Pepck1 2^nd^ (AACACTGTTTTCAAGAACACCATC & GGACATTGGGAGCCAGACT) Socs36E (GAGATCCTCACAGAGGCCACT & GCGAAACTTTCCACCTGACC) CG13220 (GCATATGCGACAAAGTGGGCC & AACATTCACCGCAAGGGCTCC) RP49 (ATCGGTTACGGATCGAACAA & GACAATCTCCTTGCGCTTCT)

For LLC and IL6-secreting LLC mouse samples, 100 mg of liver tissue was lysed in 1 mL of qiazol, using Qiagen TissueLyser II, at 20 Hz for 4 minutes. Samples were centrifuged at 12,000 rcf at 4 degrees C for 12 minutes to separate aqueous layer from organic layer and any debris. Approximately 600 microliters of RNA containing aqueous layer was collected and processed using Qiagen RNeasy lipid RNA extraction kit and QiaCube machine to isolate RNA in 50 microliter elution volumes. RNA concentrations of all samples were quantified, and samples were further diluted using ddH20 to reach a standard concentration of 100 ng/uL for each sample. TaqMan RNA-to-Ct 1 step kit was used for qPCR following provided protocol for 10 microliter reaction volumes. Data analyzed using delta delta CT method. Gapdh and Ppia were used as housekeeping gene. Primers used are Pck1 (Thermo Fisher, Mm01247058_m1), Pdk2 (Thermo Fisher, Mm00446681_m1), Pdk3 (Thermo Fisher, Mm00455220_m1), Igfbp3 (Thermo Fisher, Mm01187817_m1), Gapdh (Thermo Fisher, Mm99999915_g1), Ppia (Thermo Fisher, Mm02342430_g1).

## Protein, lipid, and carbohydrate measurements

Protocols for protein, lipid, and carbohydrate measurements were performed as previously described (Kwon et al., 2015; Ding et al., 2021). Four female flies were used for each replicate and a minimum of three replicates were measured for each sample group. Flies were homogenized in 200 ul 1X PBS with 0.1% Triton-X and Zirconium 1 mm Oxide Beads (Next Advance Lab Products, ZROB10) using TissueLyser II homogenizer (QIAGEN). Homogenate was incubated at 70°C for 10 minutes and the supernatant was collected after centrifugation at 3,000 g for 5 min. 5 ul of supernatant was applied to Pierce™ BCA Protein Assay Kit (Thermo Scientific, 23227) for detecting protein levels. TAG and free glycerol levels were quantified from 20 ul supernatant using Triglycerides Reagent (Thermo Fisher Scientific™ TR22421) and Free Glycerol Reagent (Sigma-Aldrich, F6428), respectively. Free glycerol was subtracted from TAG values. Glucose levels were measured from 10 ul supernatant using Infinity Glucose Hexokinase Reagent (Thermo Fisher Scientific™ TR15421) or D-Glucose assay kit (Megazyme, K-GLUC). Trehalose levels were measured as for glucose but incubated with 0.4 ul trehalase (Megazyme, E-TREH). The amount of glucose was subtracted from trehalose read values. TAG, free glycerol, glucose, and trehalose levels were normalized to corresponding protein levels of each sample.

## Climbing index and survival curve of flies

To assess the climbing ability, flies were transferred to a new vial and then tapped down to the bottom. Vials were imaged after 4 seconds. Percentages of flies in the upper 2/3 of the vial were recorded. 4 independent vials for each genotype were tested to generate the climbing index. Survival of flies was analyzed by calculating the percentage of flies alive in each vial incubated at 29 °C. 3 vials of flies from each genotype were tested and flies were flipped to vials with fresh food daily.

## Gut and fly imaging

Adult fly guts were dissected in cold 1X PBS and fixed for 30 minutes in 1X PBS with 4% formaldehyde. Samples were washed three times in 1X PBS with 0.3% Triton X-100 and then mounted in Vectashield with DAPI (Vector Laboratories, H-1200). Confocal images were taken with Nikon Ti and Ti2 Spinning Disk at the Microscopy Resources. Adult fly phenotypes were imaged using a ZEISS Axiozoom V16 fluorescence microscope.

## FlyPhone analysis

Cell-cell communication analysis was done using FlyPhone version 1.0 (Liu et al., 2022). Input information of ISC and EC secreted ligands and their corresponding receptor expression levels in all cell clusters were retrieved from the snRNA-seq dataset. We excluded signaling alterations that were attenuated from Day 5 to Day 8, since the wasting phenotypes were more severe at Day 8. The results are illustrated as heatmaps using TM4 software (Saeed et al., 2006).

## Chromatin Immunoprecipitation

ChIP assay was performed using SimpleChIP® Plus Enzymatic Chromatin IP Kit (Cell Signaling, 9005). For each immunoprecipitation, fat bodies from 50 adult flies with HA-tagged dominant-active Stat92e fat body expression were dissected and flash-frozen in liquid nitrogen. Samples were cross-linked with 1.5% formaldehyde for 20 minutes at room temperature (RT). After stopping cross-linking by adding glycine solution for 5 minutes at RT, samples were washed twice with 1 ml 1X PBS containing 1X Protease Inhibitor Cocktail (PIC) and disaggregated using 1 ml dounce homogenizer. Nuclei were prepared according to the manufacturer’s protocol and were lysed with Diagenode Bioruptor sonicator to release the cross-linked chromatin. Chromatins were diluted in 1X ChIP buffer and incubated with 10 ul HA Tag (C29F4) Rabbit mAb (Cell Signaling, 3724) or Normal Rabbit IgG (Cell Signaling, 2729) overnight at 4°C with rotation. 30 ul ChIP-Grade Protein G Magnetic Beads (Cell Signaling, 9006) were incubated with each immunoprecipitation for 2 hours at 4 °C with rotation. Beads were washed and incubated in 150 µl 1X ChIP Elution Buffer at 65 °C for 30 minutes with vortexing (1200 rpm) to elute the chromatin. Reversing cross-links was done by adding 6 µl 5M NaCl and 2 µl Proteinase K to the eluted chromatin supernatant and incubating 2 hours at 65°C. DNA was purified from each sample using Spin Columns provided by the kit. 1 ul DNA sample was used as template for qPCR to detect enrichments of certain DNA regions. qPCR of a fragment in the Sam-S gene region was used as the negative control. Primer used are Neg (Sam-S) (CACGGCGGCGGTGCATTCTC & CAGCGCTTGCAGAGACCGGC) Pepck1-P1 (CTAGAAAACGCTCTCAGCGCC & GCGCAGCTACGATGAGTTGG) Pepck1-P2 (GAACATATGAACGCAAAGTCCTCG & TGCTTTGTTCAATGAGCTCAGGC) Pepck1-P3 (CCTCTTGGAGGCTGGCACCA & GTTCCCTTGACACCCTCCAC) Pdk-P1 (CGTTCGCGTCAAAGTCGCGC & TTTCTCTTCTCCTGGTGCGCC) Pdk-P2 (CTCCTTGCTTCGAAGAAAGCGAG & GCGGTGAGAGGGAAGAGGAAG) Pdk-P3 (GTCGACTGTGCGCTAGACAG & TTGCAACAGGCGGTTGGCTG)

## Quantification and Statistical Analyses

We used GraphPad Prism for statistical analysis and generation of figures. Statistical analysis was done with the default settings of the software (∗ indicates p<0.05, ∗∗ indicates p<0.01, ∗∗∗ indicates p<0.001, ∗∗∗∗ indicates p<0.0001). Gene expression levels (qPCR) and metabolites levels (trehalose, glucose, TAG, and free glycerol) were normalized to the mean of control samples. Error bars indicate the standard deviations.

---

**Figure S1. Related to Figure 1.**
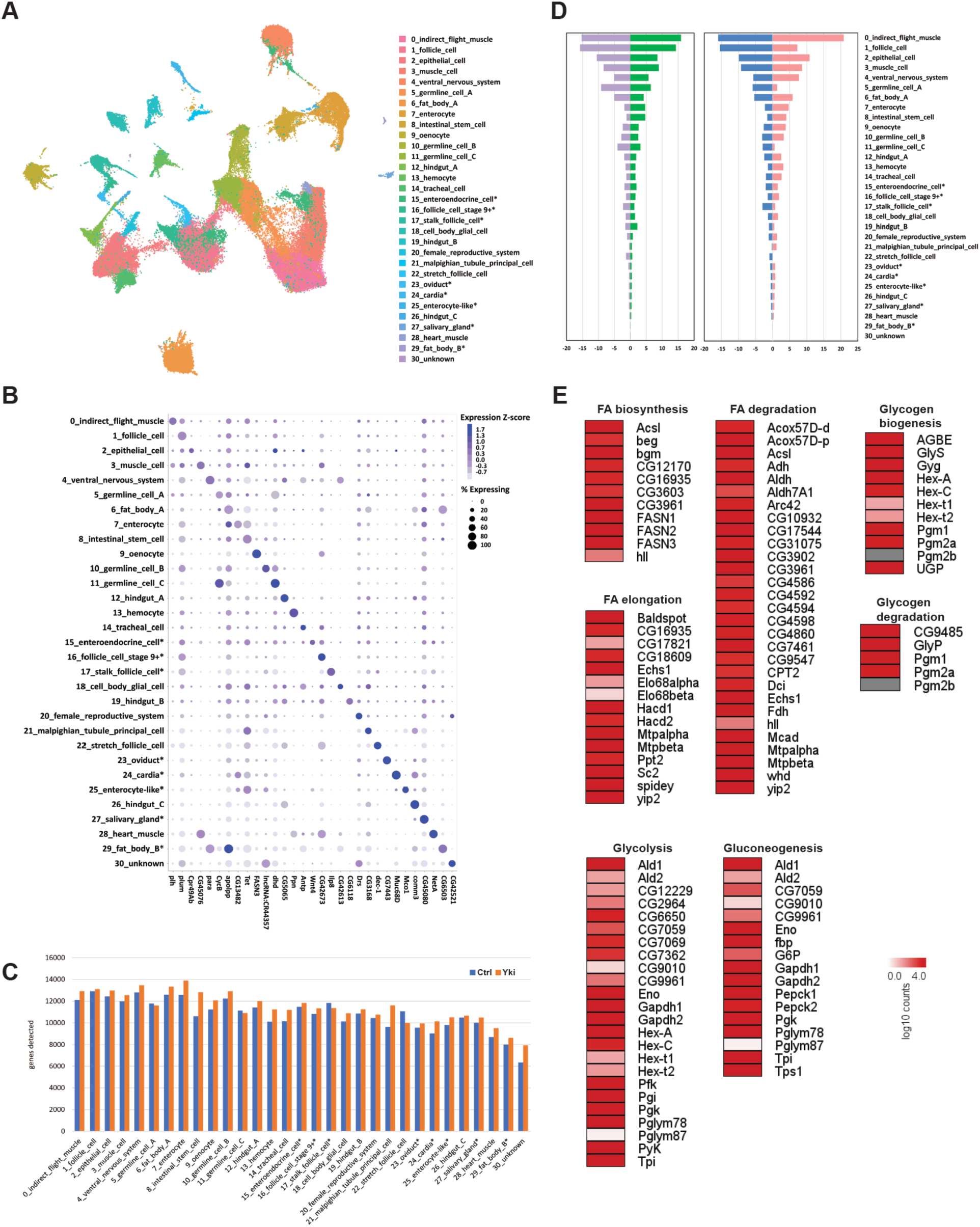
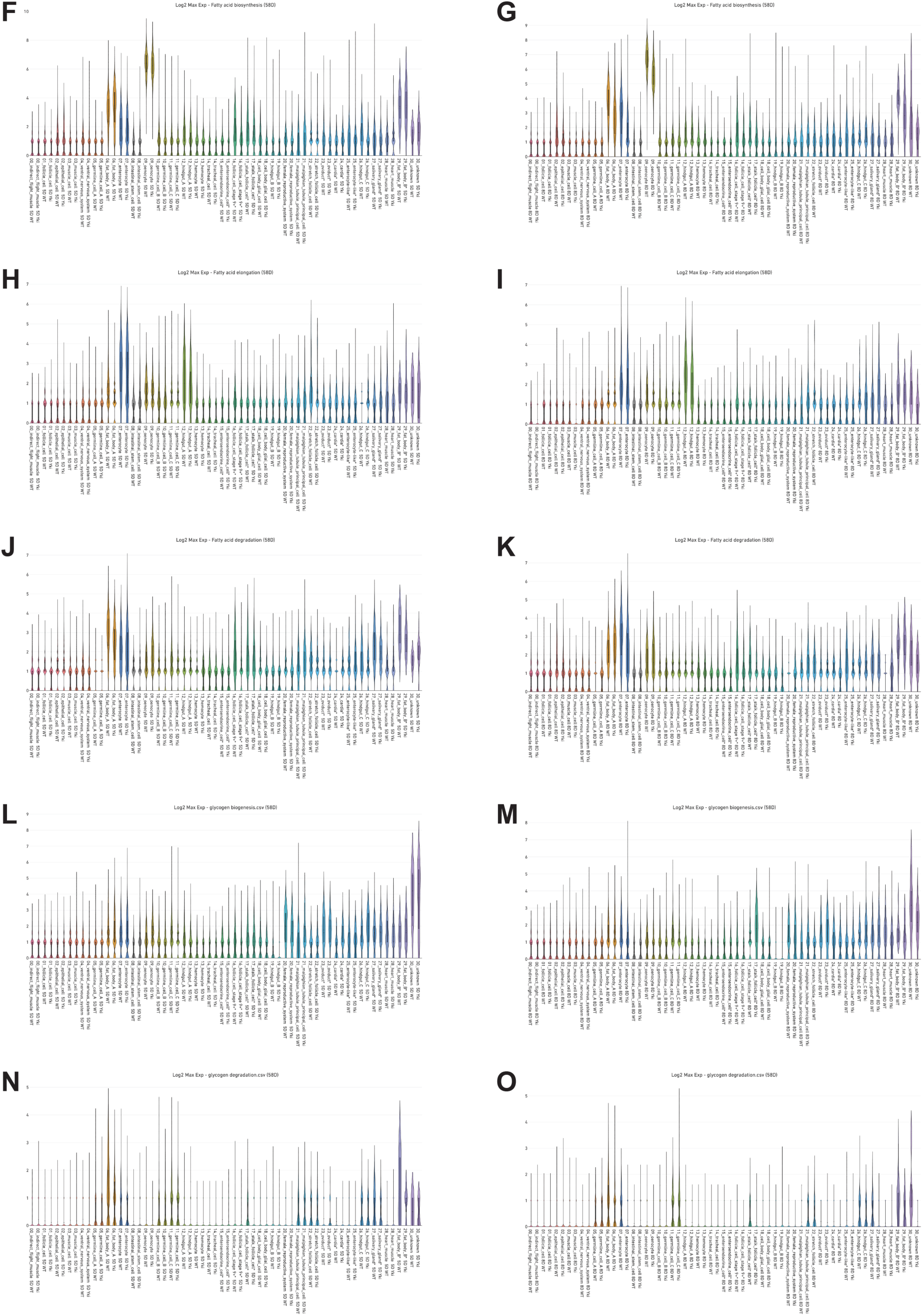
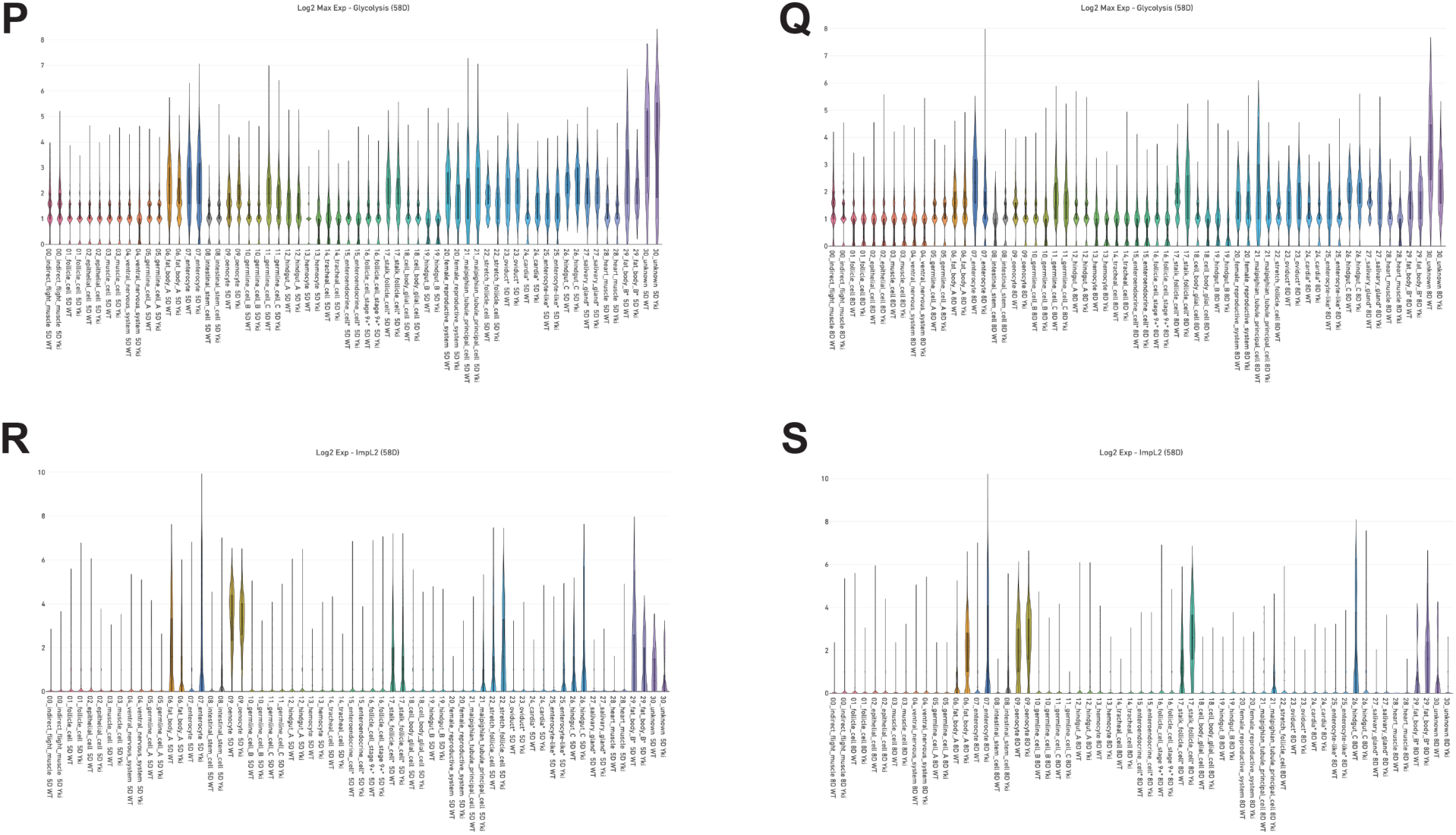
UMAP visualization of all 31 cell clusters from snRNAseq data. **(B)** Dot plot showing marker genes of each cell cluster. Color scale indicates Z-score and size indicates expression percentage. **(C)** Average genes detected in each cell cluster in the snRNAseq data. **(D)** Cell proportion comparison between control and Yki flies at Day 5 and 8. **(E)** Detection of metabolic pathway gene expression in the snRNAseq data. Expression levels of fatty acid biosynthesis pathway genes at Day 5 **(F)** and Day 8 **(G)**, fatty acid elongation pathway genes at Day 5 **(H)** and Day 8 **(I)**, fatty acid degradation pathway genes at Day 5 **(J)** and Day 8 **(K)**, glycogen biosynthesis pathway genes at Day 5 **(L)** and Day 8 **(M)**, glycogen degradation pathway genes at Day 5 **(N)** and Day 8 **(O)**, glycolysis pathway genes at Day 5 **(P)** and Day 8 **(Q)**, *ImpL2* at Day 5 **(R)** and Day 8 **(S)** in all cell clusters represented by violin plots.

**Figure S2. Related to Figure 2.**
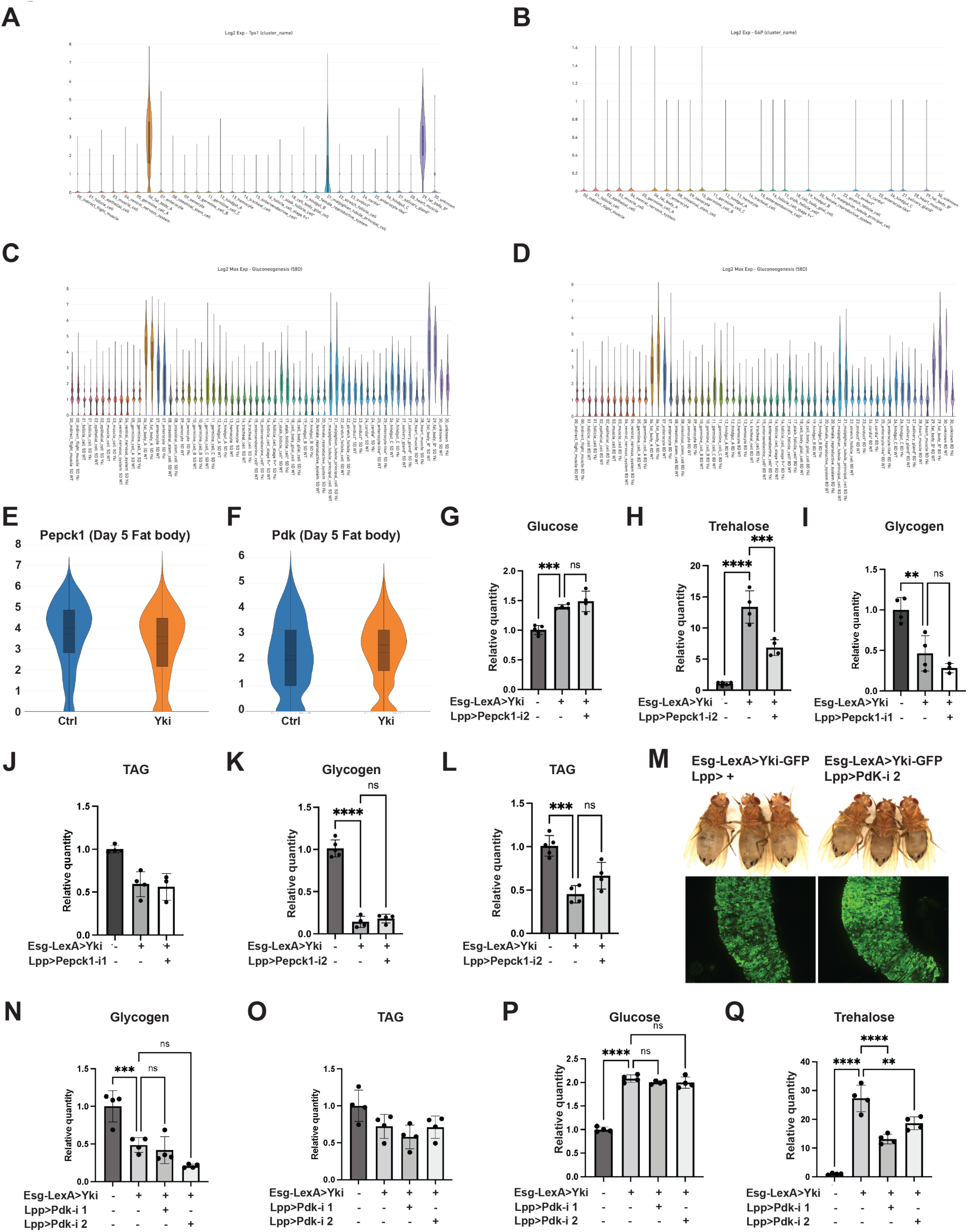
Expression levels of *Tps1* **(A)**, *G6p* **(B)**, gluconeogenesis pathway genes at Day 5 **(C)** and Day 8 **(D)** in all cell clusters represented by violin plots. Expression levels of *Pepck1* **(E)** and *Pdk* **(F)** in fat body cells at Day 5 represented by violin plot. Relative whole-body glucose **(G)**, trehalose **(H)**, glycogen **(I&K)**, and TAG **(J&L)** levels of control flies, tumor flies, and tumor flies with fat body *Pepck1* depletion at Day 8. **(M)** Representative gut tumor and phenotypes of Yki flies without and with fat body *Pdk* depletion at Day 6. Relative whole-body glycogen **(N)**, TAG **(O)**, glucose **(P)**, and trehalose **(Q)** levels of control flies, Yki flies, and Yki flies with fat body *Pdk* depletion at Day 8. **p < 0.01, ***p < 0.001, ****p < 0.0001. Error bars indicate SDs.

**Figure S3. Related to Figure 3.**
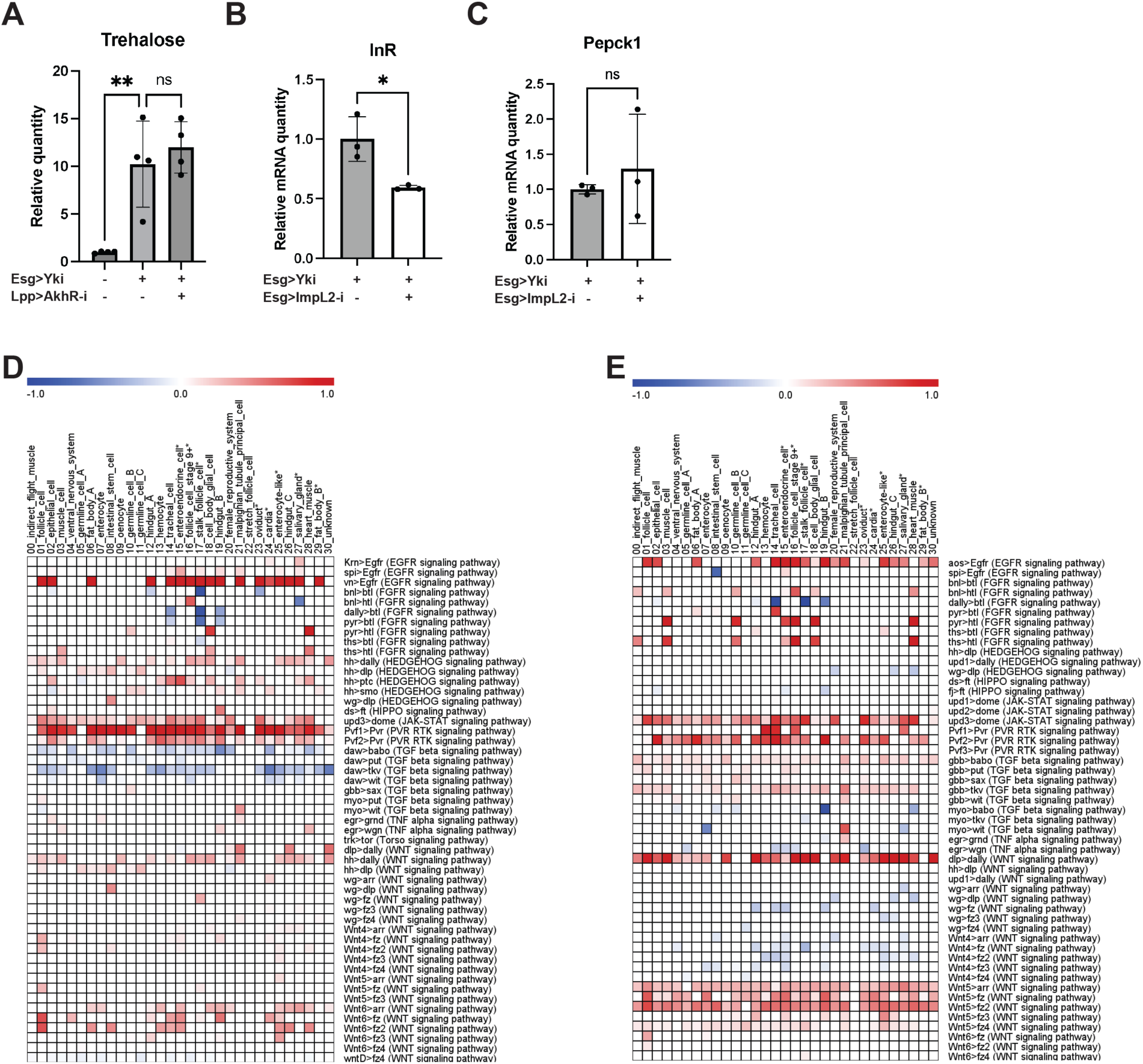
**(A)** Relative whole-body trehalose levels of control flies, Yki flies, and Yki flies with fat body *AkhR* depletion at Day 6. qRT-PCR analysis of *InR* **(B)** and *Pepck1* **(C)** mRNA levels in the fat body of Yki flies without or with ISC *ImpL2* depletion at Day 8. Heatmap plots showing perturbed signaling from EC **(D)** and from ISC **(E)** upon tumor progression (Day 8 vs Day 5). *p < 0.05, **p < 0.01. Error bars indicate SDs.

**Figure S4. Related to Figure 4.**
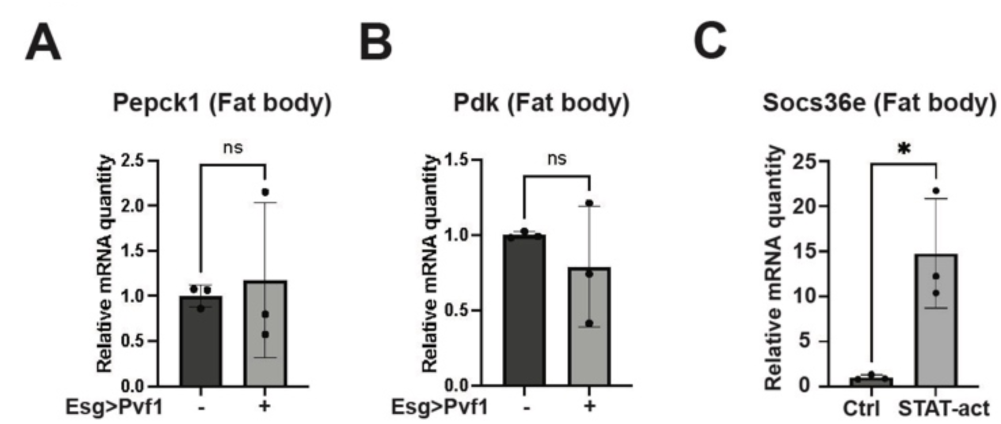
qRT-PCR analysis of *Pepck1* **(A)** and *Pdk* **(B)** mRNA levels in fat body of flies without and with ISC *Pvf1* expression at Day 8. **(C)** qRT-PCR analysis of *Socs36e* mRNA levels in fat body of flies without or with fat body *STAT-act* overexpression at Day 8. *p < 0.05. Error bars indicate SDs.

**Figure S5. Related to Figure 5.**
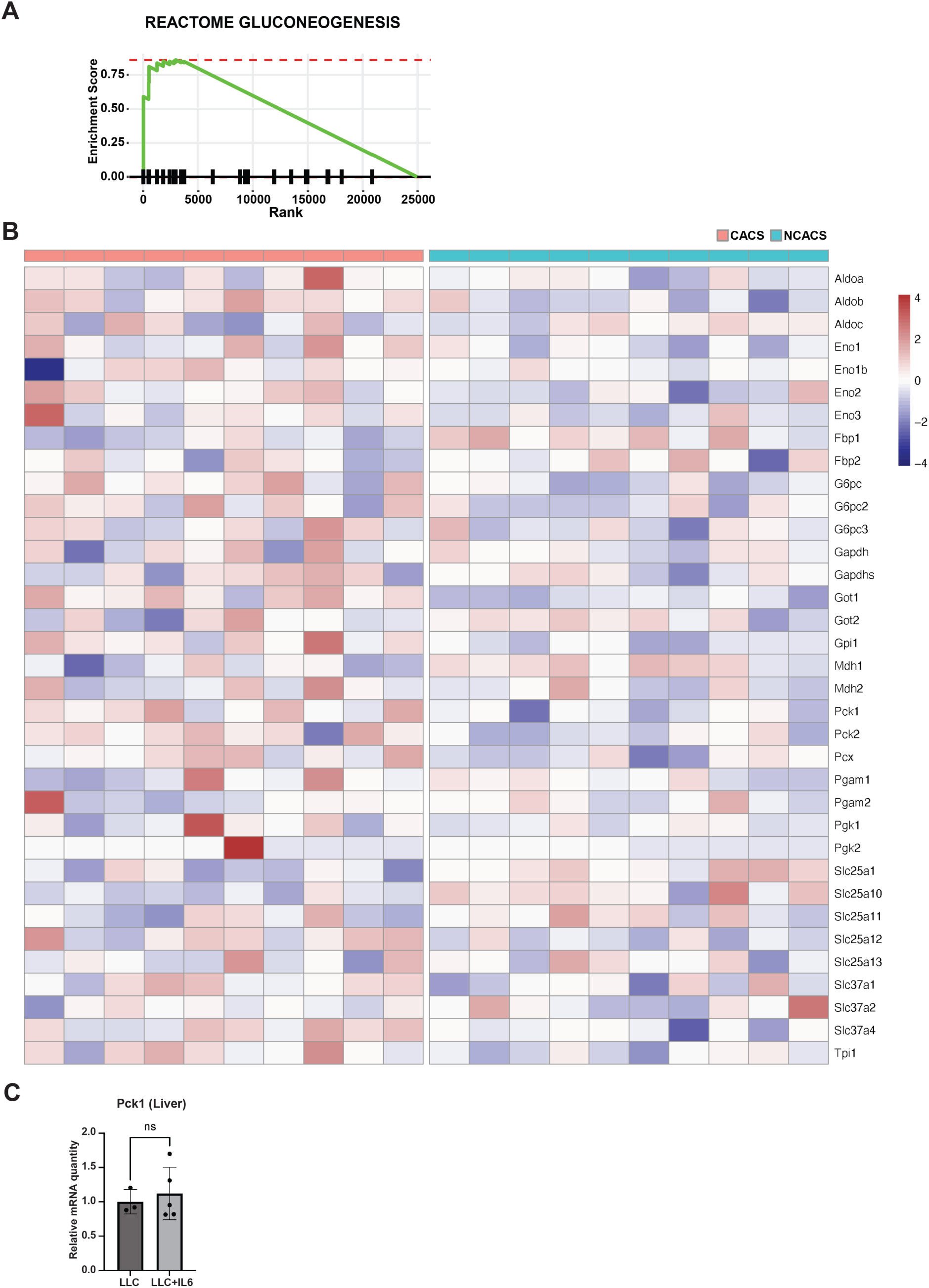
**(A)** Gene set enrichment analysis of the gluconeogenesis pathway comparing the livers of CACS to NCACS. **(B)** Heatmap of the genes from the Reactome of gluconeogenesis pathway. Columns are showing individual animals. (**C**) qRT-PCR analysis of liver *Pck1* mRNA levels of B6 mice injected with LLC cells without and with IL-6 expression.

**Figure S6. Related to Figure 6.**
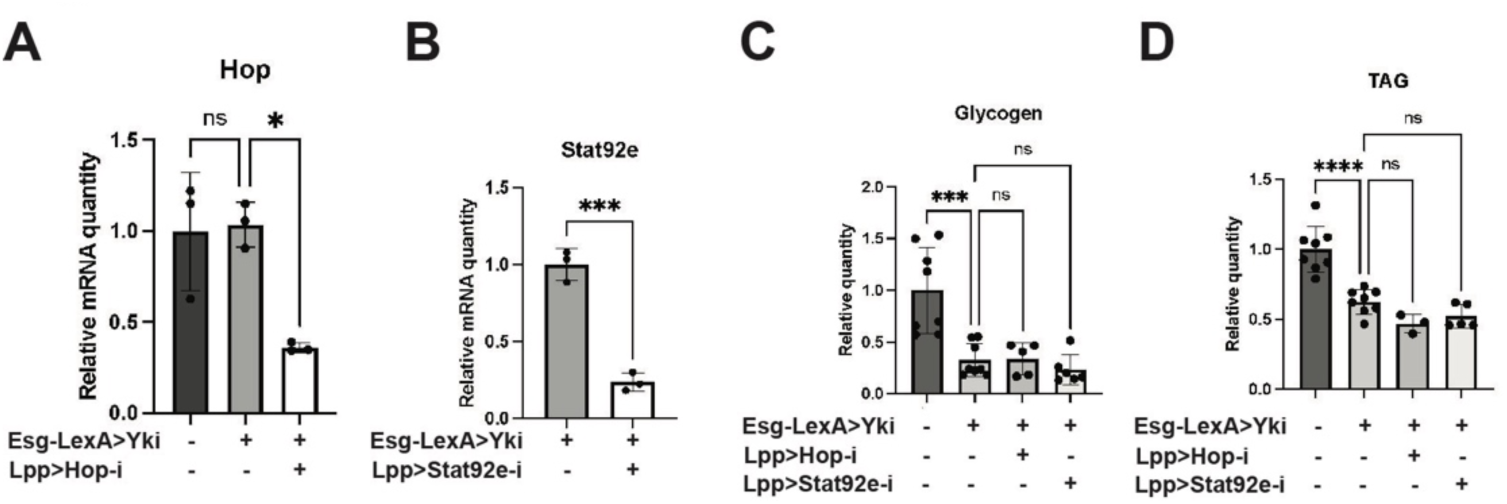
**(A)** qRT-PCR analysis of *hop* mRNA levels in fat body in control flies, Yki flies, and Yki flies with fat body *hop* depletion at Day 8. **(B)** qRT-PCR analysis of *Stat92e* mRNA levels in fat body in Yki flies without or with fat body *Stat92e* depletion at Day 8. Relative whole-body glycogen **(C)** and TAG **(D)** levels of control flies, Yki flies, and Yki flies with fat body *hop* or *Stat92e* depletion at Day 6. *p < 0.05, ***p < 0.001, ****p < 0.0001. Error bars indicate SDs.

**Table S1.**
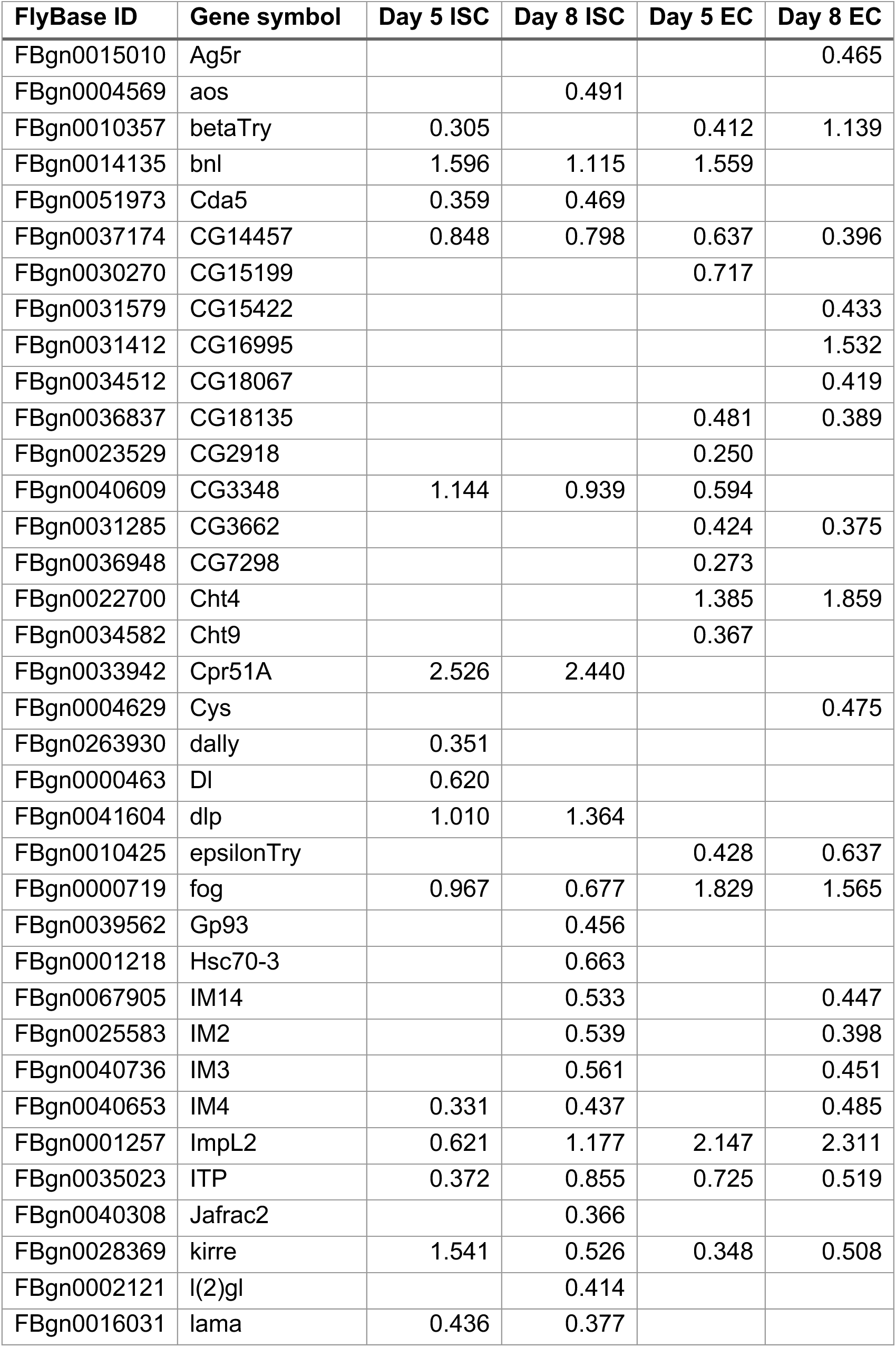
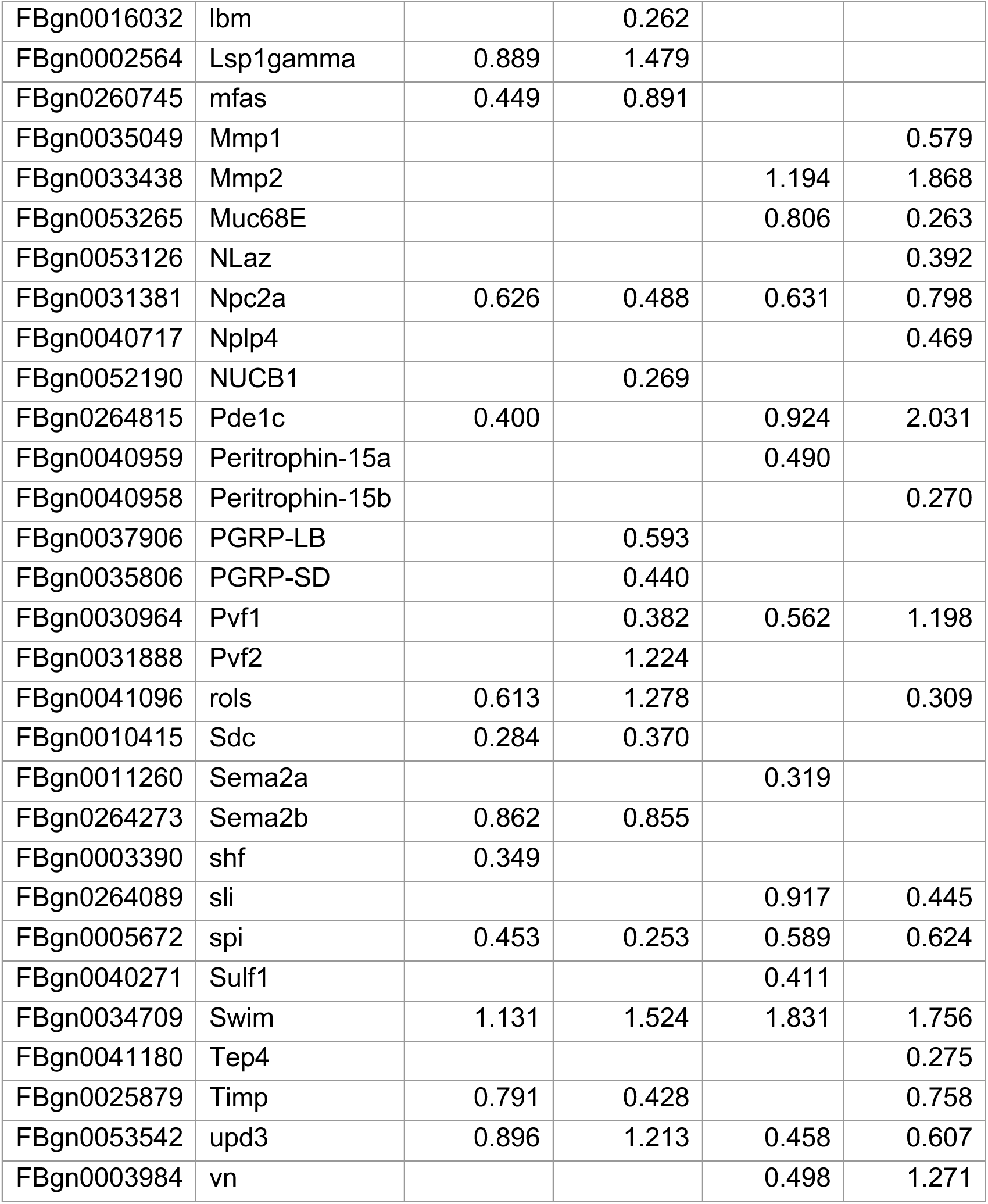
Secreted proteins upregulated in gut tumors (avg_logFC)

